# Integrative Epigenomic and High-Throughput Functional Enhancer Profiling Reveals Determinants of Enhancer Heterogeneity in Gastric Cancer

**DOI:** 10.1101/2021.06.09.447637

**Authors:** Taotao Sheng, Shamaine Wei Ting Ho, Wen Fong Ooi, Chang Xu, Manjie Xing, Nisha Padmanabhan, Kie Kyon Huang, Lijia Ma, Mohana Ray, Yu Amanda Guo, Sim Ngak Leng, Chukwuemeka George Anene-Nzelu, Mei Mei Chang, Milad Razavi-Mohseni, Michael A. Beer, Roger Sik Yin Foo, Angie Lay Keng Tan, Xuewen Ong, Anders Jacobsen Skanderup, Kevin P. White, Sudhakar Jha, Patrick Tan

## Abstract

**Background:** Enhancers are distal *cis*-regulatory elements required for cell-specific gene expression and cell fate determination. In cancer, enhancer variation has been proposed as a major cause of inter-patient heterogeneity – however, most predicted enhancer regions remain to be functionally tested.

**Results:** Analyzing 128 epigenomic histone modification profiles of primary GC samples, normal gastric tissues, and GC cell lines, we report a comprehensive catalog of 75,730 recurrent predicted enhancers, the majority of which are tumor-associated *in vivo* (>50,000) and associated with lower somatic mutation rates inferred by whole-genome sequencing. Applying Capture-based Self-Transcribing Active Regulatory Region sequencing (CapSTARR-seq) to the enhancer catalog, we observed significant correlations between CapSTARR-seq functional activity and H3K27ac/H3K4me1 levels. Super-enhancer regions exhibited increased CapSTARR-seq signals compared to regular enhancers even when decoupled from native chromatin contexture. We show that combining histone modification and CapSTARR-seq functional enhancer data improves the prediction of enhancer-promoter interactions and pinpointing of germline single nucleotide polymorphisms (SNPs), somatic copy number alterations (SCNAs), and *trans*-acting TFs involved in GC expression. Specifically, we identified cancer-relevant genes (e.g. *ING1*, *ARL4C*) whose expression between patients is influenced by enhancer differences in genomic copy number and germline SNPs, and HNF4α as a master *trans*-acting factor associated with GC enhancer heterogeneity.

**Conclusions:** Our study indicates that combining histone modification and functional assay data may provide a more accurate metric to assess enhancer activity than either platform individually, and provides insights into the relative contribution of genetic (*cis*) and regulatory (*trans*) mechanisms to GC enhancer functional heterogeneity.

## Introduction

Enhancers are a specific class of *cis*-regulatory elements that play important roles in regulating cell-specific gene expression [1, 2]. Enhancer dysregulation has been reported to contribute to multiple human diseases, including complex conditions such as cancer, Alzheimer’s disease, and diabetes [3, 4]. Specific to cancer, recent studies have highlighted enhancer dysregulation as a pervasive feature of malignancy [5–7], and the functional impact of enhancer heterogeneity, both between and within tumors, in the establishment and maintenance of tumour phenotypes, cancer prognosis, and treatment response [6, 8, 9]. For example, transcriptome-defined subtypes of medulloblastoma have distinct enhancer landscapes and cellular origins [5]. Moreover, genomic alterations including single nucleotide polymorphisms (SNPs), somatic mutations, chromosomal rearrangements, and somatic copy number alterations (SCNAs) can drive enhancer-linked phenotypic heterogeneity [10–14]. Examples include focal amplifications of super-enhancers near the *MYC* locus in 17% of lung adenocarcinomas [11], and elevated *FOXA1* expression and genomic occupancy at specific enhancers driving metastases in subgroups of pancreatic cancer [12]. In ovarian cancer, the rs7874043 germline SNP within a *PSIP1*-linked enhancer affects Sp1 binding, and the *C* allele is associated with poor progression-free survival [13].

Super-enhancers comprise a cluster of putative enhancers occurring in close genomic proximity, typically characterized by high levels of transcription factor (TF), Mediator (MED) binding, and active chromatin marks such as histone H3 lysine 27 acetylation (H3K27ac) [15, 16]. Relative to typical enhancers, super-enhancers have been reported to be more significantly associated with tumour-specific gene expression, cancer hallmarks and disease-associated genetic variation [15, 17, 18]. However, conflicting data persists in the field as to whether super-enhancers truly exhibit distinct functional characteristics and properties compared to regular enhancers, or whether they are simply assemblies of regular enhancers [19, 20]. One possible reason for this controversy is that the field’s ability to accurately identify functional enhancers and to assess their contribution to target gene expression (ideally in a quantitative manner) remains challenging. While H3K27ac enrichment has been widely utilized as a surrogate of enhancer activity, predicting enhancers based on histone ChIP data can result in both false-positive and false-negative findings [21], and assigning specific target gene(s) to distal enhancers remains an active area of research [22]. Traditionally, reporter assays such as luciferase assays have been used to directly quantify enhancer strength [23]; however, there are currently very few data sets in the public domain where such reporter measurements are available in a high-throughput scale. More recently, highly parallelized enhancer functional assays, such as STARR-seq and CapSTARR-seq, have been described as high-throughput and quantitative approaches to assess enhancer activity in mice, humans and cancer cells, facilitating the identification of enhancers and enhancer-gene relationships [24–26].

Gastric cancer (GC) is one of the most common cancer worldwide and a leading cause of global cancer-associated mortality, showing high incidence in East Asia, East Europe, Central and South America [27]. Individual gastric tumors are highly heterogeneous with different subtypes and distinct molecular characteristics, clinical outcomes, and responses to therapy [28]. Recent studies from our group and others have highlighted a role for epigenetic alterations and enhancer heterogeneity in GC [29]. Here, by integrating enhancer information from both CapSTARR-seq and histone-ChIPseq, we explored the mechanistic basis of enhancer heterogeneity in GC.

Combining CapSTARR-seq and histone profiling data, we identified germline SNPs, SCNAs, and *trans*-acting transcription factor driving enhancer heterogeneity between different GC patients. We also provide evidence that combining histone modification and CapSTARR-seq functional data may provide a more accurate metric to assess enhancer activity than either platform individually.

## Results

### Predicted enhancer landscapes in GC cell lines reflect regulatory function *in vivo*

We analyzed 128 chromatin profiles covering multiple histone H3 modifications (H3K27ac, H3K4me1 and H3K4me3) across 28 GC cell lines, 4 gastric normal cells, 18 primary GCs and 18 matched normal gastric tissues. Of these profiles, approximately one-third (44 profiles) were newly generated for this study, and have not been previously reported. Supplementary Table 1,2 provides clinical information and sequencing statistics. Several of the profiles were generated using Nano-ChIP-seq, previously shown to exhibit good concordance with both conventional ChIP-seq and also orthogonal ChIP-qPCR results[29, 30]. We performed two independent quality control assessments of the Nano-ChIP-seq data: ChIP-seq enrichment analysis over known promoters, and assessment by the quality control software CHANCE (CHip-seq ANalytics and Confidence Estimation) [31]. Comparison of input-corrected ChIP-seq and input signals at 1000 promoters associated with highly expressed protein-coding genes demonstrated successful enrichment in the vast majority (99%) of H3K27ac and (100%) H3K4me3 libraries (Supplementary Table 2). CHANCE analysis also revealed that the majority (77%) of ChIP libraries exhibited successful enrichment, including H3K4me1. Besides Nano-ChIP-seq, the samples were also characterized by DNA methylation profiling, ATAC-seq (Assay for Transposase-Accessible Chromatin using sequencing), whole-genome sequencing (WGS), and Illumina RNA-sequencing. To map candidate *cis*-regulatory elements on a genome-wide scale, we identified genomic regions of H3K27ac enrichment, previously shown to be associated with active promoters and enhancers [32] (Fig. 1a). We initially focused on the GC cell lines, to identify enhancers active in epithelial cancer cells and to avoid potential stromal contamination. Retaining H3K27ac ChIP libraries with >13,000 CCAT-detected peaks, we selected peaks distal from annotated TSS sites (>2.5k), excluding four GC lines (NCC24, NUCG4, RERF-GC-1B and YCC22). Using this approach, we identified 5,451-111,571 predicted distal enhancers per GC cell line. The wide range of predicted enhancers across GC lines is unlikely explained by technical variation, as they are highly reproducible across biological replicates (R= 0.86-0.89, Fig. 1b and Supplementary Fig. 1). Our results are also similar to previous studies of luminal breast cancer, where the number of predicted enhancers across biological samples ranges from 2,383-172,092

**Figure 1.**
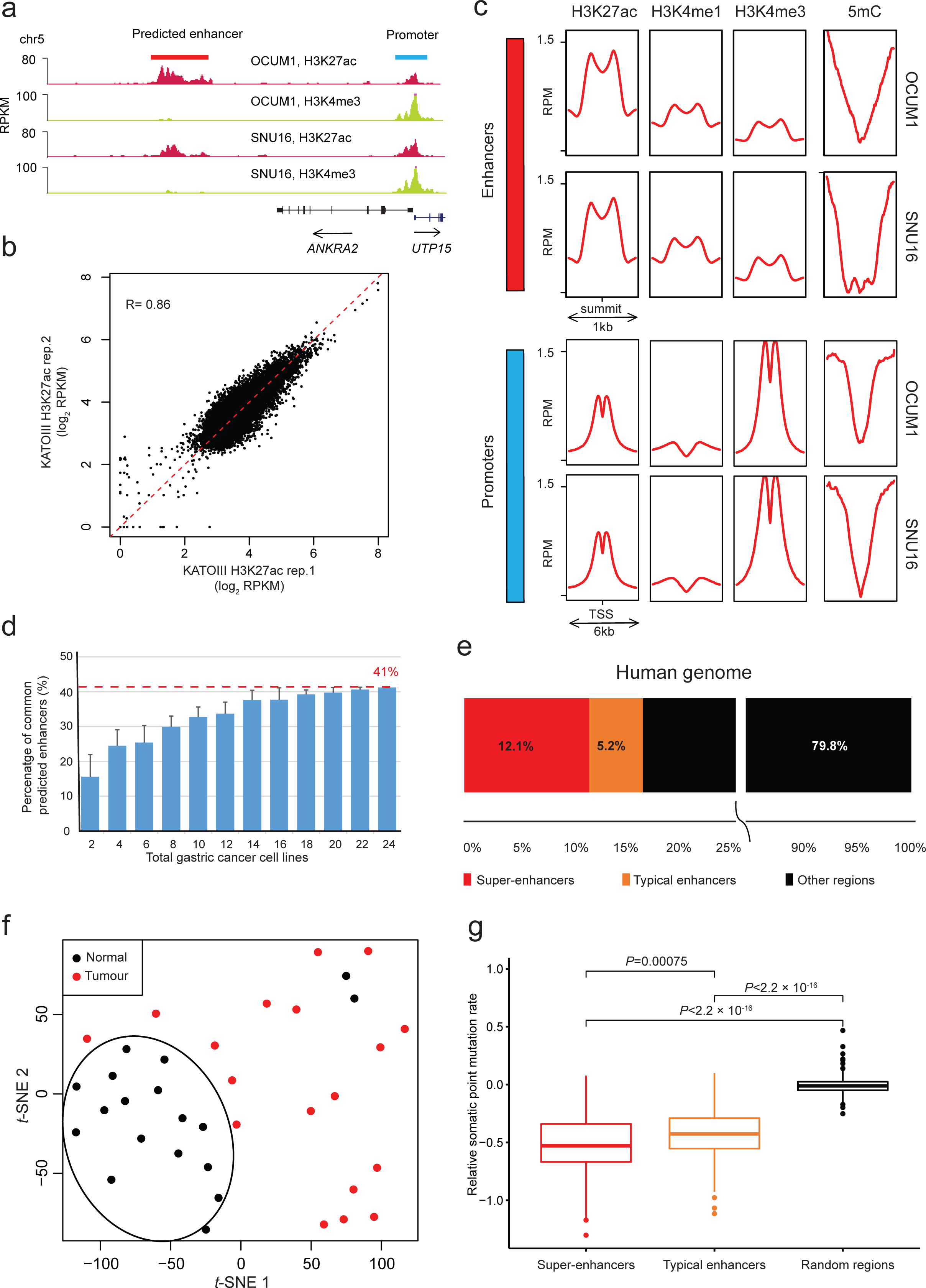
Distal enhancer landscapes of GC cell lines. **a)** Histone profiles of OCUM1 and SNU16 cells show enrichment of H3K27ac and H3K4me3 around the *UTP15* TSS. A predicted distal enhancer enriched for H3K27ac and >2.5kb distant from *UTP15* TSS is observed. **b)** Comparison of H3K27ac signals over common predicted enhancers between two KATOIII replicates. **c)** Genome-wide profiling of chromatin marks (H3K27ac, H3K4me1, and H3K4me3) and DNA methylation (5mC) at predicted enhancers and promoters. **d)** Recurrence rates of predicted enhancers. Data presented are the mean percentage +/-standard deviation of common predicted enhancers found in two or more gastric cancer cell lines, as a function of number of cell lines. **e)** Distribution of predicted super-enhancer and typical enhancers present in GCs across the human genome. **f)** *t*-distributed stochastic neighbour embedding (*t*-SNE) analysis using predicted enhancers present in GCs reveals separation between GCs and matched normal tissues (n =36). **g)** Difference in somatic point mutation rates among predicted super-enhancers, typical enhancers, and randomly selected genomic regions. *P*-value: two-side Student’s t-test.

[6].

Concordant with previous studies [25, 32, 33], the predicted enhancer regions manifested enriched bimodal H3K27ac signals, H3K4me3 signal depletion, H3K4me1 signal enrichment, and were distinct from promoter-associated regions that are H3K4me3/H3K4me1 positive (Fig. 1c). We detected 183,788 predicted distal enhancer regions in total, where some enhancers occurred in multiple lines (“recurrent”; present in 2 or more lines) while other regions existed in only one line (“private”). The frequency of recurrent predicted enhancers approached saturation at about 18 GC lines, larger than the number of GC lines analyzed in previous studies (n=11; Fig. 1d) [29].

Increasing the recurrence threshold from 2 to 3 did not appreciably alter the number of cell lines required for saturation (Supplementary Fig. 2). We thus defined a comprehensive GC enhancer catalog of ∼75K enhancers present in 2 or more lines (n = 75,730).

Using the ROSE algorithm [34], we also identified super-enhancers from the GC enhancer catalog, corresponding to enhancer subgroups spanning large genomic regions and associated with high levels of transcription factor (TF) binding and H3K27ac histone marks. We identified 483 to 2,089 predicted super-enhancers per GC line and a consensus set of 8,293 super-enhancers across the lines. Supporting previous findings that super-enhancers play important roles in cell identity and cancer hallmarks, we observed super-enhancers associated with several known GC oncogenes, such as *MYC*, *KLF5* and *EGFR* (Supplementary Fig. 3).

To determine which cell-line-predicted enhancers are also present *in vivo*, we compared H3K27ac enrichment levels for these regions across 18 primary GCs. Of 75,730 cell-line-predicted enhancers, around two-thirds (n = 52,457) were also present in two and more primary GC samples (Fig. 1e). Supporting their association with cancer malignancy, *t*-Distributed Stochastic Neighbor Embedding (*t*-SNE [35]) using the predicted enhancers confirmed separations between GCs and matched normal tissues (Fig. 1f). In addition, integrating somatic mutation rates from previously published WGS data from 212 GCs confirmed that regions harboring predicted super-enhancers/enhancers were associated with significantly lower somatic mutation rates (*P*<2 × 10^-16^, Student’s t-test, Fig. 1g). As *cis*-regulatory elements are known to be more accessible to DNA repair complexes, this result further supports the *in vivo* validity of the GC enhancer catalog.

### Functional enhancer profiling reveals intrinsic regulatory potential and elevated super-enhancer activity

While H3K27ac-enrichment is widely used as an epigenetic surrogate of enhancer activity, only a limited number of studies have experimentally investigated the extent to which H3K27ac-enriched regions exhibit *bona-fide* transcriptional activity. To explore if enhancer regions predicted by H3K27ac can indeed function as true enhancers, we employed CapSTARR-seq technology [24], a high-throughput approach for quantitatively assessing the activities of thousands of enhancers. Briefly, in CapSTARR-seq, DNA fragments bearing candidate enhancer elements are captured using custom-designed probes and cloned by homologous recombination into mammalian vectors downstream of a basal promoter. The CapSTARR-seq library is then transfected *en masse* into cell lines, and RNA-seq is performed to detect mRNA reads corresponding to the enhancer sequence (Fig. 2a). Notably, because enhancer elements in CapSTARR-seq are cloned into exogenous plasmids, CapSTARR-seq transcriptional signals are likely to reflect intrinsic transcriptional activity of the enhancer, in the absence of other factors such as larger-scale chromatin patterns and topologically associating domains (TADs).

**Figure 2.**
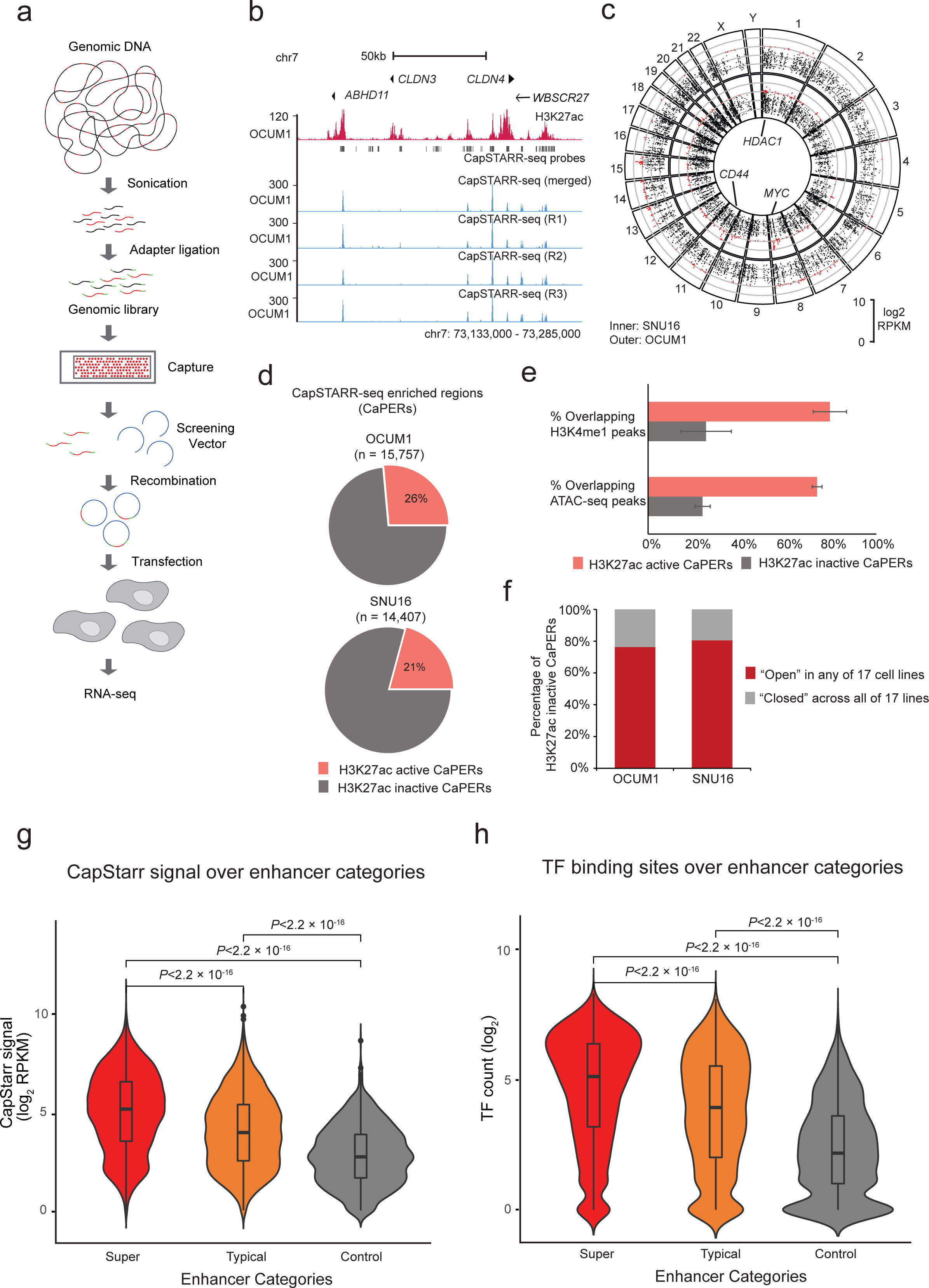
CapSTARR-seq Functional Enhancer Profiling. **a)** CapSTARR-seq experimental workflow. **b)** H3K27ac (red) and CapSTARR-seq (blue) profiles at the *ABHD11*, *CLDN3* and *CLDN4* loci in OCUM1 cells. The top blue track depicts CapSTARR-seq signals averaged from three CapSTARR-seq replicates. Black boxes denote CapSTARR-seq probes. **c)** Circos visualization of CapSTARR-seq signals across the human genome in SNU16 (the inner circle) and OCUM1 cells (the outer circle). *HDAC1*, *MYC* and *CD44* are highlighted as genes associated with CapSTARR-seq high enhancers. **d)** Distribution of H3K27ac active and inactive elements in CapSTARR-seq enriched regions (CaPERs) in OUCM1 and SNU16 cells. **e)** Differences in H3K4me1 ChIP-seq and ATAC-seq peak enrichment at H3K27ac active and inactive CaPERs. Error bars indicate two independent biological replicates. **f)** Distribution of OCUM1/SNU16 H3K27ac inactive CaPERs in the “open” state across 17 cell lines. Red regions denote OCUM1/SNU16 H3K27ac inactive CaPERs located in open chromatin in any of 17 cell lines. Grey regions denote OCUM1/SNU16 H3K27ac inactive CaPERs located in closed chromatin across all the cell lines. **g)** Differences in CapSTARR-seq signals (log2 RPKM) in enhancer categories in OCUM1 cells. *P*-values are calculated using the Mann–Whitney U test. **h)** Differences in TF binding sites over enhancer categories. The number of TF binding sites over an enhancer was calculated using the ReMap database. *P*-values are calculated using the Mann–Whitney U test.

Informed by the GC enhancer catalog, we designed CapSTARR-seq probes targeting 78,974 regions (median size 5 kb). In brief, 57.6% of the CapSTARR-seq probes (n=45,526) were located within super-enhancers; 16.3% within typical enhancers (n=12,863); and as negative controls we included 20,585 probes (n=20,585; 26.1%) capturing genomic regions outside of predicted enhancer regions. The CapSTARR-seq library, covering ∼100,000 genomic regions, was transfected into two GC lines (SNU16 and OCUM1), and RNA-seq was performed to provide a quantitative assessment of enhancer activity. We selected OCUM1 and SNU16 cells as cell line models for two reasons. First, OCUM1 and SNU16 cells were validated as two lines that closely resemble GC tumors by Celligner, a tool aligning tumor and cell line transcriptional profiles [36]. Second, OCUM1 and SNU16 have been previously used as GC models in many other published studies, and are thus widely regarded as accepted GC models in the field [29, 37]. CapSTARR-seq enriched regions were identified using MACS2 (q value < 0.05). We observed high correlations between CapSTARR-seq replicates performed in GC cell OCUM1, indicative of the high reproducibility of the assay (Fig. 2b). Fig. 2c depicts CapSTARR-seq activities of enhancers captured across the whole human genome in OCUM1 or SNU16 cells.

We defined CapSTARR-seq functional enhancer activity as the CapSTARR-seq fold change (FC) over input signal and demarcated three groups based on FC values: weak (1.5 ≤ FC < 2), moderate (2 ≤ FC < 3) and strong (FC ≥ 3). We then proceeded to explore relationships between levels of epigenetic mark enrichment (e.g. H3K27ac) against CapSTARR-seq activity. As anticipated, enhancers exhibited significantly higher CapSTARR-seq signals compared to negative controls (P<2 × 10^-16^, Student’s t-test, Supplementary Fig. 4). When analyzed against H3K27ac, H3K4me1, H3K4me3 and DNA methylation, we found that strong enhancers displayed the highest H3K27ac and H3K4me1 signals while moderate enhancers were associated with intermediate levels (H3K27ac: P<0.01; H3K4me1: P<0.05; Supplementary Fig. 5). All three CapSTARR-seq categories were also depleted for 5-methylcytosine (5mC), a marker of gene repression and inactivation (Supplementary Fig. 5).

Although in aggregate enhancer regions exhibited significant consistency between CapSTARR-seq activity and H3K27ac signals, in absolute terms only 21 – 26% of CapSTARR-seq enriched regions (CaPERs) overlapped H3K27ac-enriched region in the lines tested (Fig. 2d). As a representative example, Supplementary Fig. 6 depicts a H3K27ac inactive CaPER at the *SNX30* genomic region in OCUM1 cells, exhibiting high CapSTARR-seq signals but H3K27ac and ATAC-seq peak depletion. We reasoned that these results might be due, at least in part, by CapSTARR-seq measuring signals from cytoplasmic reporter constructs and thereby likely reflecting a region’s *potential* regulatory activity independent of *native* chromatin context. Two findings support this model. First, in SNU16 and OCUM1 cells, we found that regions with positive CapSTARR-seq signals but inactive by H3K27ac were associated with reduced H3K4me1 and ATAC-seq signals, implying that these are ‘dormant’ enhancers with lower accessibility to *trans*-acting factors (Fig. 2e). Second, among CapSTARR-seq enriched regions, approximately four-fifths of enhancers “dormant” in either OCUM1 or SNU16 cells exhibited an “open” chromatin state (i.e. active ATAC-seq signals) in at least one of 17 GC lines (Fig. 2f). These findings thus suggest that CapSTARR-seq and H3K27ac data are complementary, with the former reflecting a regions regulatory potential, while the latter reflects native chromatin accessibility.

We next asked if super-enhancers and typical enhancers might exhibit distinct patterns of CapSTARR-seq activity. We found that super-enhancer regions demonstrated statistically higher functional enhancer signals than typical enhancers (*P*<2.22 × 10^-16^, Mann–Whitney U test, Fig. 2g). Using a generalized linear model (GLM), we confirmed that the increased activity levels of super-enhancers were independent of differences in DNA accessibility (assessed by ATAC-seq), DNA copy number levels, and genomic length (Supplementary Fig. 7). When analyzed at the level of DNA sequence and comparing TF binding sites over each predicted enhancer (using the Remap database [38]), we discovered that transcription factor (TF) binding occupancy was associated with increasing functional enhancer strength (Fig. 2h). These results demonstrate that even when decoupled from their original chromatin context, regions associated with super-enhancers still exhibit higher functional enhancer activity than typical enhancers, and this increased activity is likely due to an abundance of dense TF *cis*-binding occupancy at super-enhancer regions.

### Combining CapSTARR-seq enhancer activity with H3K27ac profiling may improve prediction of enhancer-promoter connections

Due to their localization at distal genomic regions, accurately assigning specific genes to enhancers (“enhancer-gene connections”) remains challenging [39, 40]. Previous studies have used H3K27ac-enrichment as a surrogate of enhancer activity to predict enhancer-promoter connections [41, 42]. To explore if CapSTARR-seq functional enhancer activity might enhance the prediction of enhancer-promoter interactions, either alone or combined with H3K27ac, we employed the recently-published Activity-by-contact (ABC) model [43], previously developed to predict functional enhancer–gene connections. Briefly, in the ABC model, the quantitative effect of a regulatory element on the expression of a target gene depends on its own strength as an enhancer (Activity) weighted by the frequency of its 3D contact with the gene’s promoter (Contact). The relative contribution (ABC score) of an enhancer to one gene’s expression is calculated by that enhancer’s effect divided by the total effect of all enhancers associated with the gene (Fig. 3a).

**Figure 3.**
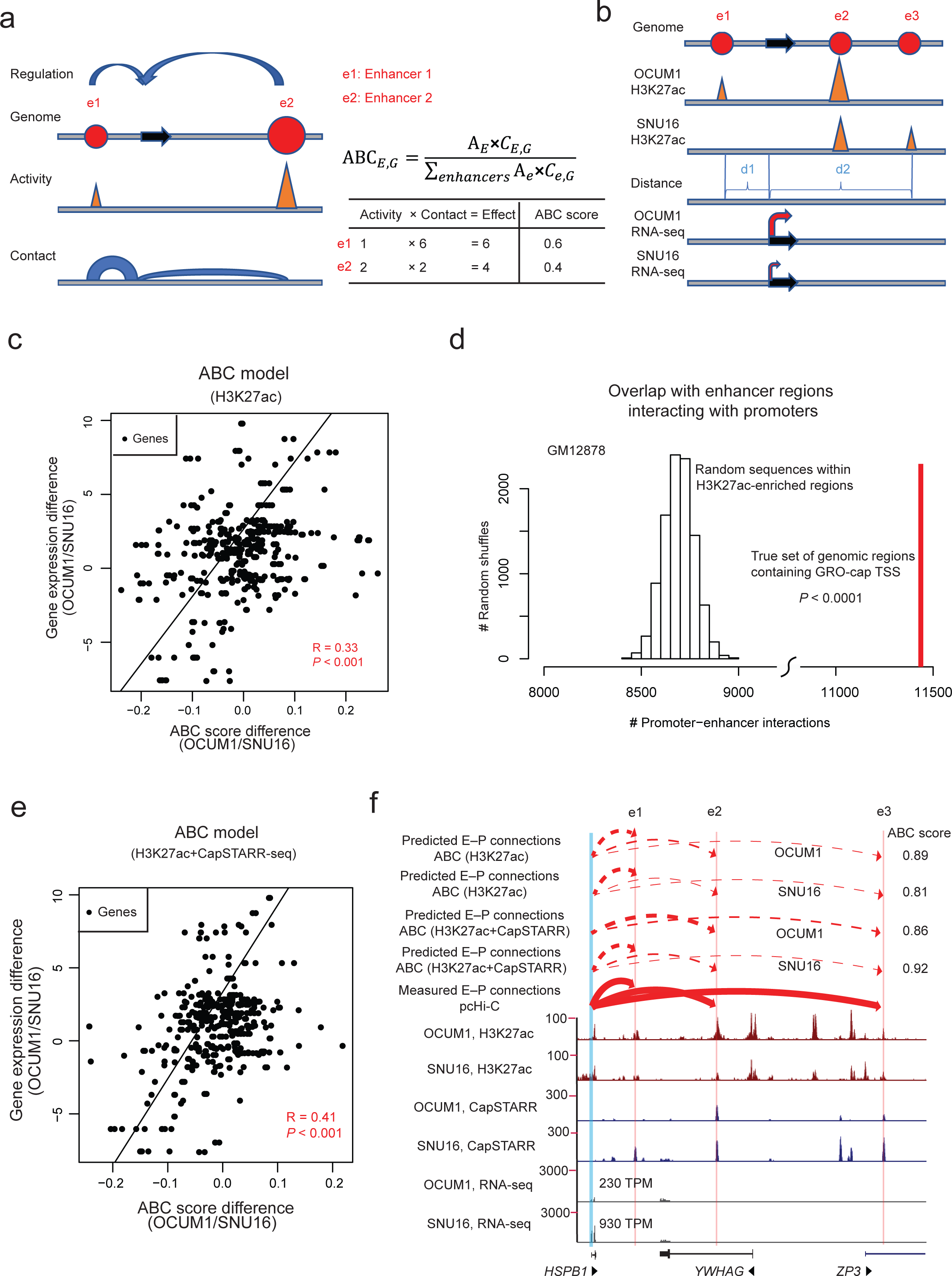
Activity-by-contact model of enhancer-gene regulation. **a)** ABC model schema. e1 and e2 denote two arbitrary enhancers (solid red circles) for a gene (black arrow). Activity estimates the enhancer strength while Contact estimates the frequency of the enhancer-gene connection. The ABC score of an enhancer to one gene’s expression is calculated by that enhancer’s effect divided by the total effect of all enhancers for the gene. **b)** ABC model for explaining gene expression differences between OCUM1 and SNU16 cells. Activity is estimated as the level of H3K27ac-enrichment at an enhancer region while Contact is quantified as a function of the genomic distance between the enhancer and the TSS of the gene (Contact = Distance^-1^). Gene expression level is estimated from RNA-seq data. **c)** Comparison of ABC score differences and observed gene expression differences between OCUM1 and SNU16 cells. Each dot represents a differentially expressed gene between OCUM1 and SNU16 cells. Activity of an enhancer is estimated as the H3K27ac signal. R: Pearson’s correlation coefficient. *P*-value: Pearson’s correlation test. **d)** Genomic regions containing GRO-cap TSS overlap more enhancer regions interacting with promoters than randomly shuffled regions in GM12878 lymphoblastoid cells. *P*-value calculated empirically by random shuffling of sequences within H3K27ac-enriched regions. **e)** Comparison of ABC score differences and observed gene expression differences between OCUM1 and SNU16 cells. Activity of an enhancer is estimated as the geometric mean of H3K27ac-enrichment and CapSTARR-seq signal. R: Pearson’s correlation coefficient. *P*-value: Pearson’s correlation test. **f)** Comparison of the *HSPB1* expression difference and the ABC score difference between OCUM1 and SNU16 cells in the *HSPB1* locus (chr7:75,927,230-76,048,225). Predicted E–P connections (dotted red arcs) are based on ABC maps in OCUM1 and SNU16 cells. Observed E–P connections (solid red arcs) are derived from the pcHi-C database.

We first examined if differential enhancer landscapes based on differences in H3K27ac ChIP-seq signals can explain (at least in part) gene expression differences between OCUM1 and SNU16 cells using the ABC model. Operationally, we estimated Activity as the level of H3K27ac-enrichment at an enhancer region, and Contact as a function of the genomic distance between the enhancer and the TSS of the gene (Contact = Distance^-1^) (Fig. 3b). We selected 430 highly differentially expressed genes between OCUM1 and SNU16 cells. For each gene, we included all H3K27ac-predicted enhancers within 5 Mb of the gene’s promoter as candidate functional enhancers. The total effect of all functional enhancers on gene expression was estimated as the sum of ABC scores of those enhancers. We found that gene expression differences between OCUM1 and SNU16 were significantly correlated with ABC score difference (Pearson R= 0.33*, P* = 1.44 × 10^-12^, Fig. 3c).

Recent studies have reported that transcription occurring at distal enhancers (“enhancer transcription”) is predictive of enhancer function, with potentially higher accuracy than histone modifications [44]. To explore if enhancer transcription can also improve identification of enhancer-promoter interactions, we first analyzed publicly available GRO-cap TSS-positive enhancer regions interacting with promoters in GM12878 lymphoblastoid cells. Of 34,436 GRO-cap TSS-positive enhancers, we found 11,439 cases (33%) involved in enhancer-promotor interactions. These interactions were independently validated by promoter capture Hi-C (pcHi-C) data [45] and were higher than enhancer-promoter connections inferred using H3K27ac alone (∼8,693 expected by chance inside H3K27ac active regions, *P*<0.0001 vs. random shuffling of sequences within H3K27ac active regions, Fig. 3d), indicating that enhancer transcription may be a more specific predictor of enhancer function than H3K27ac signals. Next, to ask if native enhancer transcription can be inferred from cytoplasmic STARR-seq data, we reanalyzed “High-resolution Dissection of Regulatory Activity” (HiDRA) datasets [46], which test putative regulatory regions by coupling accessible chromatin extraction with self-transcribing episomal reporters (ATAC-STARR-seq). From 7,000,000 DNA fragments in GM12878 lymphoblastoid cells, we identified ∼47,000 enhancer regions enriched for HiDRA signals referred to as “STARR active enhancers”. In parallel, by reanalyzing H3K27ac ChIP-seq profiles of the same cell line, we identified ∼ 61,000 enhancer regions showing significant H3K27ac enrichment, referred to as “H3K27ac active enhancers”. Within the H3K27ac active enhancers, 44% of STARR active elements exhibited native enhancer transcription (as determined by GRO-cap TSS profiling). By contrast, only approximately 9% of STARR inactive elements within the H3K27ac active enhancers were GRO-Cap TSS positive (Supplementary Fig. 8; *P* < 2.2 × 10^-16^, two-sided Fisher’s exact test.). We validated these GRO-seq, STARR-seq and H3K27ac relationships in a separate cell line (K562; Supplementary Fig. 8). These results suggest that STARR active enhancers are more likely to exhibit native enhancer transcription and involved in enhancer-promoter interactions.

We thus tested if incorporating CapSTARR-seq functional enhancer activity also enhances the performance of the ABC model in our GC data, estimating Activity as the geometric mean of H3K27ac-enrichment and CapSTARR-seq signals at an enhancer region, rather than H3K27ac alone. We observed a significant and improved correlation between gene expression difference and ABC score difference (Pearson R= 0.41, *P* < 2.2 × 10^-16^, Fig. 3e). The ABC model incorporating H3K27ac-enrichment and functional enhancer activity outperformed that using either of the estimators individually (H3K27ac only: Pearson R = 0.33, *P* =0.05; CapSTARR-seq only: Pearson R = 0.20, *P* <0.001, Dunn and Clark’s Z test [47]). For example, an ABC model based on H3K27ac estimated enhancer activity only connected three enhancers to the *HSPB1* promoter in both OCUM1 and SNU16 cells (Fig. 3f). These three enhancer-promoter interactions were validated by the pcHi-C database. Although the model predicted that the sum of ABC scores (SNU16: 0.81; OCUM1: 0.89) of these three enhancers was larger in OCUM1 cells than SNU16 cells, *HSPB1* exhibited higher expression in SNU16 cells compared to OCUM1 cells (SNU16: 930 TPM; OCUM1: 230 TPM). However, by adding Cap-STARRseq enhancer activity into the ABC model, the *HSPB1* expression difference between OCUM1 and SNU16 cells became consistent with the ABC score difference. We reasoned that the activation of the enhancer e1, which exhibited CapSTARR-seq activity, may increase *HSPB1* expression in SNU16 cells. By contrast, enhancer e1 exhibited no CapSTARR-seq activity in OCUM1 cells. Taken collectively, these results indicate that incorporating Cap-STARRseq enhancer activity may elevate the accuracy of estimating enhancer functionality and the ability of predicting enhancer-gene connections when coupled with H3K27ac ChIP-seq data.

### Effects of large-scale copy number alterations on differential enhancer activity

We then combined the CapSTARR-seq and H3K27ac acetylation patterns (see Methods) to explore mechanisms associated with enhancer heterogeneity in GC. In total, we identified 1,888 differential enhancers between OCUM1 and SNU16 cells. We hypothesized that differential enhancers might result from at least two sources – *cis*-based genomic variation (CNVs and SNPs), and *trans*-based TF binding (Supplementary Fig. 9 provides a cartoon comparing two samples). In the ‘*cis*-model’, sample-specific SCNAs or SNPs result in differential enhancer activity, perhaps by influencing the recruitment of TFs commonly expressed in both samples. In the ‘*trans*-model’, differential expression of TFs between samples may underlie sample-specific alterations in enhancer activity.

To test the influence of SCNAs on differential enhancer activity, we applied CNVkit [48] on WGS data of OCUM1 and SNU16 to infer SCNAs in both lines. When summarized across all differential enhancers, we observed that H3K27ac and CapSTARR-seq signals were positively correlated with SCNAs (*P* < 4.6 × 10^-13^, Supplementary Fig. 10). Of the 1,888 differential enhancers, approximately one quarter (513 enhancers, 27.2%) exhibited concordance between H3K27ac signals and DNA copy numbers in OUCM1 and SNU16 cells (Fig. 4a). To extend these results, we then queried 26 additional cell lines, and identified 58 enhancers showing significant associations between H3K27ac and SCNA levels, of which 35 also showed a statistically significant correlation with target gene expression levels. These included known oncogenes such as *FGFR2* and *MYC* (Supplementary Table 3). Notably, we observed a significant correlation between copy number deletions of an *ING1*-associated enhancer, decreased H3K27ac signals, and decreased *ING1* expression in multiple lines (Fig. 4b). *ING1* encodes a protein that physically interacts with the TP53 tumor suppressor and negatively regulates cell growth [49]. It has been reported that *ING1* may function as a tumor suppressor gene in gastric cancer [50]. Using overexpression vectors, we confirmed that overexpression of *ING1* in GC cells significantly reduced cell proliferation (Supplementary Fig. 11), consistent with a purported anti-oncogenic function.

**Figure 4.**
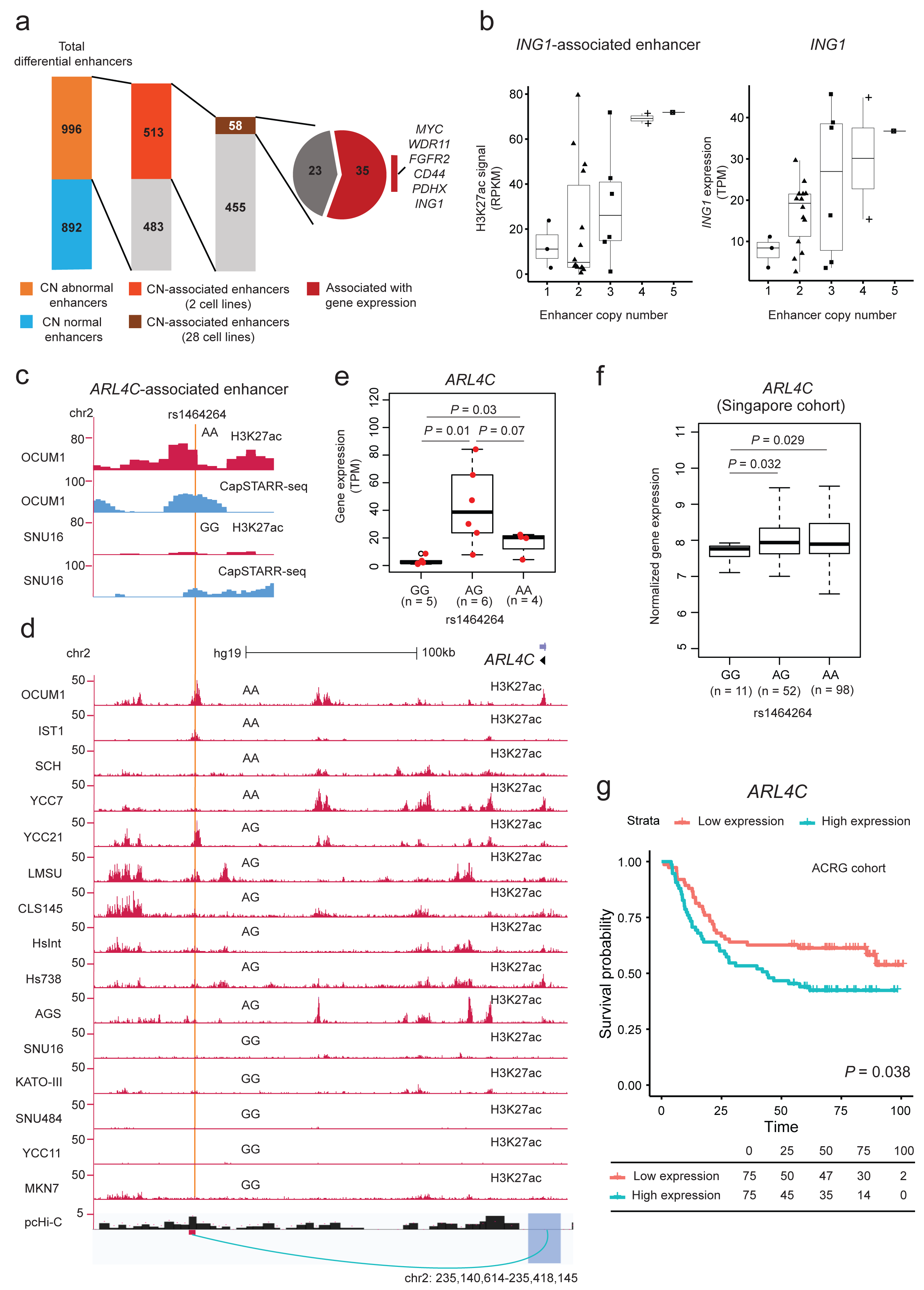
*Cis*-analysis of differential enhancers. **a)** Distribution of enhancers associated with copy number. CN-abnormal enhancers represent differential enhancers with altered DNA copy numbers in OCUM1 or SNU16. CN-associated enhancers represent differential enhancers showing concordant changes in H3K27ac signals and DNA copy number. **b)** Correlation of H3K27ac signals and DNA copy numbers for the *ING1* enhancer and with *ING1* gene expression. **c)** H3K27ac ChIP-seq and CapSTARR-seq tracks at the enhancer region harboring the *ARL4C*-associated SNP rs1464264 in OCUM1 and SNU16 cells. **d)** H3K27ac ChIP-seq tracks in GC lines and pcHi-C associations in gastric tissues. The SNP rs1464264 genotype is annotated above the H3K27ac ChIP-seq track in each cell line. Significant chromatin interactions are shown below the axis (green loop). **e)** Differences in expression of *ARL4C* in three groups of cell lines with different SNP rs1464264 genotypes (GG, AG and AA). *P*-value: Mann–Whitney U test. **f)** Difference in expression of *ARL4C* in three groups of GC patients with different SNP rs1464264 genotypes (GG, AG and AA) in the Singapore cohort (n= 161). *P*-value: Student’s t-test. **g)** Survival analysis comparing patient groups with samples exhibiting low (green) and high (red) expression of *ARL4C* in the ACRG cohort (n=300). *P*-value is calculated using the Log-rank test. Survival data are indicated for every 25 months.

### Effects of germline SNP variation on differential enhancer activity

Next, we proceeded to explore the role of germline SNP variation in differential enhancer activity. Here, we restricted our analysis to differential enhancers located in diploid regions in both lines (“Copy number-normal enhancers”), excluding the confounding effects of SCNAs. Within 892 copy number-normal enhancers, we identified 605 unique germline SNPs using Genome Analysis Toolkit (GATK) between OCUM1 and SNU16 cells. To further discover SNPs whose genotype correlates with histone acetylation levels, we applied the genotype-independent signal correlation and imbalance algorithm (G-SCI [51]) across a cohort of 28 cell lines. Specifically, we identified SNPs correlating with cohort-variation in H3K27ac-enrichment over the same regions, referred to as histone acetylation QTLs (haQTLs). We identified 207 haQTLs (34%) using the G-SCI test. Interestingly, this discovery rate is significantly higher than (8%) reported in previous studies [51] despite both studies using the same controlled false-discovery rate (FDR) of 0.1. It is likely that the higher haQTL discovery rate in our study might be due to the combination of CapSTARR-seq and H3K27ac ChIP-seq data, that allows finer-scale resolution of enhancer boundaries. Supporting this, we also applied G-SCI to differential enhancers only based on H3K27ac. In brief, of 8,016 SNPs within H3K27ac-defined differential enhancers, 2,194 HaQTLs (27%) were detected using the G-SCI test (FDR < 0.1). Two-side proportion test gave us a significant difference when comparing the two ratios (*P* =3.2 × 10^-4^).

Of the 207 haQTLs, we focused on 43 germline SNPs predicted by RegulomeDB [52], a database of human regulatory variants reported to influence protein DNA binding (RegulomeDB score 1 or 2, Supplementary Table 4). For instance, haQTL rs1464264 is located at an OCUM1-specific enhancer (Fig 4c), and histone acetylation levels over this enhancer region was associated with the rs1464264 genotype among multiple cell lines (Fig. 4d). Specifically, increased H3K27ac at enhancers covered by the major *A* allele coincided with increased expression of *ARL4C* (ADP-ribosylation factor-like 4C, a class of GTP-binding protein), suggesting that the *A* allele potentially upregulates the gene (Fig. 4e). Moreover, the enhancer region harboring rs1464264 exhibits long-range interactions with the *ARL4C* gene promoter region as evidenced by pcHi-C database (*P*<10^-3^, Fig. 4d). The correlation of the *A* allele and increased *ARL4C* expression was also confirmed in an independent validation set of primary GC patients from Singapore (n=161 patients, *P*=0.029 between GG and AA, Fig. 4f). *ARL4C* was significantly upregulated in GC compared to normal samples in the ACRG cohort (*P*=0.008; Supplementary Fig. 12). Survival analysis revealed patients with GCs exhibiting high expression of *ARL4C* showed poor overall survival compared with GC samples where *ARL4C* is relatively lowly expressed in the ACRG cohort (*P* = 0.038, Log-rank test, Fig. 4g). It has been reported that *ARL4C* is a peritoneal dissemination-associated gene in GC [53]. We confirmed that knockdown of *ARL4C* in GC cells using two independent shRNAs significantly reduced cell proliferation (Supplementary Fig. 13), consistent with a purported oncogenic function.

### Effects of *trans*-acting TF binding on differential enhancer activity

Finally, we explored the role of *trans*-acting factors in differential enhancer activity. Using HOMER, a *de novo* motif discovery algorithm, we found that OCUM1-specific enhancers (n=1,014) were enriched in FRA1, FRA2, and ATF3, while SNU16-specific enhancers (n= 847) exhibited enrichments in FOXA1, HNF4α and KLF5/6 binding (Fig. 5a, Supplementary Fig. 14). Although OCUM1-specific enhancers defined only based on H3K27ac were also enriched in FRA1, FRA2, and ATF3 binding, the significance of enrichment and the percentage of target sequence with motif were lower, indicating that combination of CapSTARR-seq and H3K27ac ChIP-seq data may enable stronger enrichment of TFs. In brief, the average rate of target sequence with FRA1, FRA2, and ATF3 motif on OCUM1-specific enhancers defined based on two signals was twice as much as that on enhancers defined only based on H3K27ac (32% vs 14%, Supplementary Fig. 14). Among these TFs binding on SNU16-specific enhancers, we focused on HNF4α, as HNF4α was the only TF exhibiting significantly upregulated expression in GCs as compared to normal tissues in two independent GC cohorts (Singapore cohort: *P* = 4.9 × 10^-3^; TCGA cohort: *P* = 1.8 × 10^-4^; Mann– Whitney U test, Fig. 5b).

**Figure 5.**
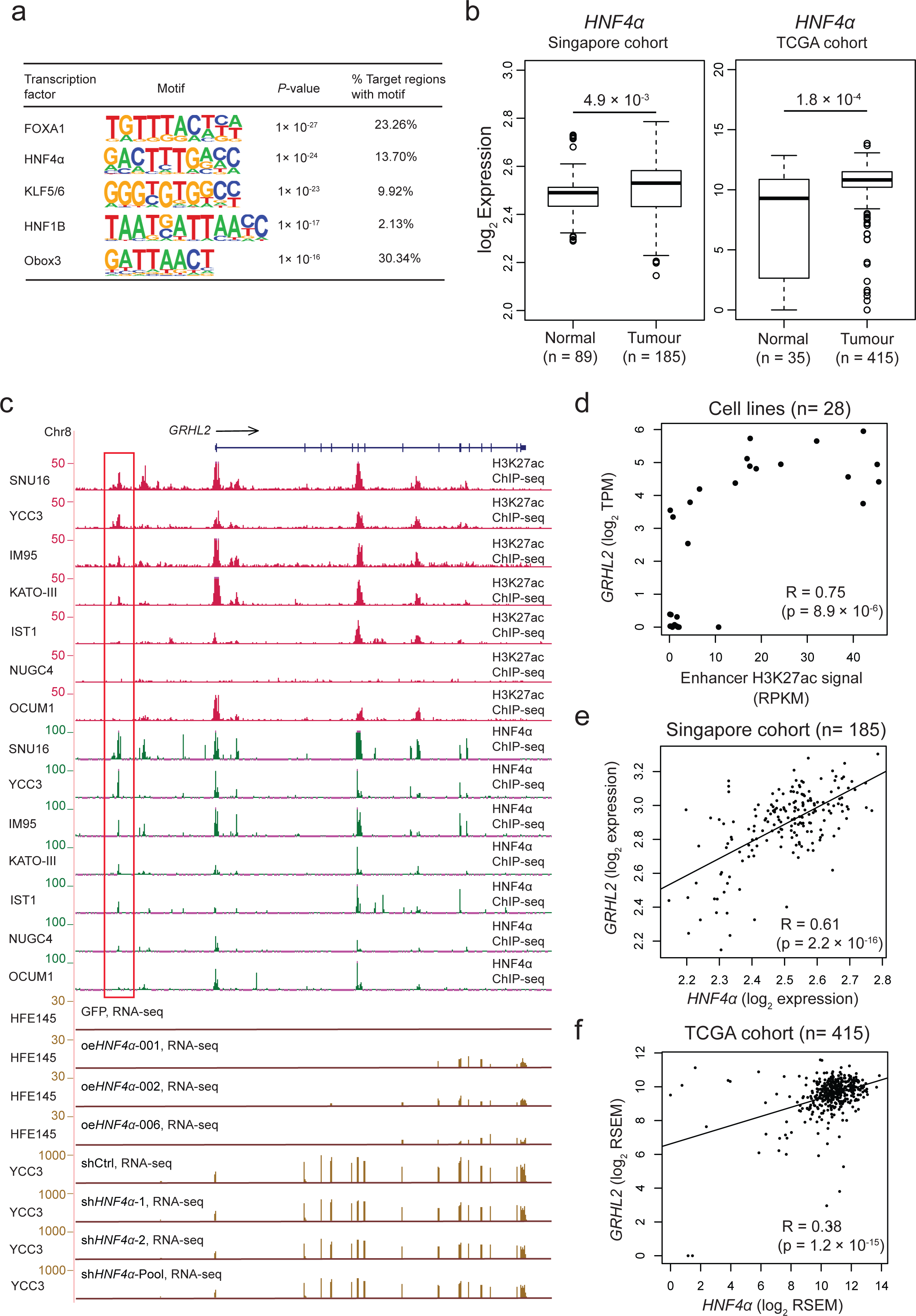
*Trans*-analysis of differential enhancers. **a)** Top 5 transcription factor binding enrichments at SNU16-specific enhancers determined by HOMER *de novo* motif analysis. **b)** Expression of *HNF4α* in normal gastric (n= 89) and GC samples (n=185) from the Singapore cohort. Expression of *HNF4α* in normal gastric (n=35) and GC samples (n=415) from the TCGA cohort. *P*-value: Mann–Whitney U test. **c)** Integration of H3K27ac ChIP-seq data, HNF4α ChIP-seq binding profiles (SNU16, YCC3, IM95, KATO-III, IST1, NUGC4 and OCUM1) and RNA-seq data in control, *HNF4α* overexpressing cells (HFE145), *HNF4α* knockdown (YCC3) at the *GRHL2* gene locus. The red box indicates an enhancer associated with *GRHL2* (chr8: 102,449,130-102,450,795). **d)** *GRHL2* gene expression levels and H3K27ac signals over the enhancer for *GRHL2* are linearly correlated across 28 cell lines. R: Pearson’s correlation coefficient. *P*-value: Pearson’s correlation test. **e)** Singapore cohort analysis reveals *GRHL2* and *HNF4α*transcriptomic correlations using microarray data (Pearson’s correlation test). **f)** TCGA cohort analysis confirms *GRHL2* and *HNF4α* transcriptomic correlations using RNA-seq data (Pearson’s correlation test).

To identify differential enhancers associated with HNF4α binding, we focused on SNU16-specific enhancers exhibiting 1) occupancy of HNF4α in SNU16 cells; 2) a significant correlation between HNF4α binding enrichment and H3K27ac signals among multiple lines. Of 64 enhancers out of 847 fulfilling these criteria, 22 were further highlighted as they exhibited a statistical correlation between H3K27ac signals and the expression levels of their corresponding target genes (Supplementary Table 5). For example, Fig. 5c highlights one enhancer near the *GRHL2* locus, where *GRHL2* expression levels are significantly correlated with H3K27ac signals over the enhancer region among multiple GC lines (Pearson R = 0.75, *P* = 8.9 × 10^-6^, Fig. 5c). This region was selected for two more reasons: 1) *GRHL2* is significantly up-regulated in GCs (Supplementary Fig. 15); 2) *GRHL2* has recently been reported to be involved in Epithelial-mesenchymal transition in GC [54]. To confirm that *GRHL2* is a target gene of *HNF4a*, we performed *HNF4a* overexpression and knockdown experiments in GC lines. We observed concordant dysregulation of *GRHL2* RNA levels upon HNF4α perturbation (Fig. 5c). To independently validate this relationship *in vivo*, we further confirmed a positive correlation of *GRHL2* and *HNF4a* expression in both the Singapore cohort and the TCGA cohort (Singapore cohort: *P* = 2.2 × 10^-16^; TCGA cohort: *P* = 1.2 × 10^-15^; Pearson’s correlation test, Fig. 5e, 5f). Taken together, our results suggest that differential HNF4α binding enrichment at this enhancer is associated with differential H3K27ac ChIP-seq and CapSTARR-seq signals, and also *GRHL2* expression.

## Discussion

In this study, we surveyed the GC enhancer landscape across 28 GC cell lines, 4 normal gastric cell lines, 18 primary GCs and 18 matched normal gastric tissues. To our knowledge, this epigenomic data set currently represents the largest such data set for GC, expanding previous studies by over one-third and conceptually extending previous findings by applying CapSTARR-seq to directly test >70,000 candidate H3K27ac-positive enhancer elements. Previous studies using STARR-seq (a predecessor to CapSTARR-seq) in *Drosophila* have demonstrated that enhancer strengths correlate well with average gene expression [55], and for STARR-seq-high enhancers occurring nearby lowly expressed genes, these are explainable by the former not being accessible in their native chromatin context. In our study, we similarly identified a cadre of enhancer elements showing strong CapSTARR-seq signals but not exhibiting high H3K27ac levels in the same cell line. However, these regions exhibited H3K27ac signals when extrapolated to other cell lines, suggest that these regions are likely ‘dormant’ enhancers that can activate gene expression in the context of a hospitable and open chromatin environment. Supporting this idea, others have reported that strong STARR-seq enhancers located at closed chromatin regions can drive nearby gene expression when they are opened by TSA, a histone deacetylase (HDAC) inhibitor [25].

The question of whether super-enhancers comprise a novel paradigm that is distinct from regular enhancers remains to be determined. By analyzing the α-globin super-enhancer, Hay *et al.* reported that each constituent enhancer “acts independently and in an additive fashion” [56]. In contrast, Shin *et al.* reported that super-enhancers associated with the *Wap* gene operate across a temporal and functional hierarchy of constituent enhancers [57]. In our study, we found that even when decoupled from their chromatin context, genomic regions associated with super-enhancers exhibit higher functional enhancer activities compared to regular enhancers. Our results suggest that this increased enhancer activity is likely due to an intrinsic abundance of dense TF *cis*-binding occupancy sites. Other studies have also reported that individual enhancers within super-enhancers can exhibit stronger activating features, such as H3K27ac-enrichment and TF binding densities [15–17]. Therapeutically, regions exhibiting dense TF binding occupancies have been shown to represent important regulatory nodes susceptible to changes in TF concentration [17]. Such vulnerabilities to TF perturbation at super-enhancers may offer productive targets for cancer therapy. It will be interesting to explore this area further using other methods of functional enhancer testing, such as recently developed CRISPR interference (CRISPRi) and CRISPR activation (CRISPRa) platforms [58, 59].

Identifying enhancer-promoter interactions remains challenging because promoters and regulatory elements can be separated by millions of base pairs [60], and oftentimes the closest gene is usually not the true enhancer target [22, 43, 61]. Correlation-based approaches identify enhancer-promoter connections by combining genomic proximity with genetic associations (eg expression quantitative trait loci (eQTLs)) or chromatin state, enabling detection of long-range interactions [62, 63]. Here, we found that combining CapSTARR-seq enhancer activity with H3K27ac profiling improves the prediction of gene expression differences. Previous studies have shown that incorporating STARR-seq activity can enable fine-scale resolution of active enhancer boundaries derived from lower-resolution chromatin immunoprecipitation (ChIP) profiles [44, 64]. Supporting these results, we found that combining CapSTARR-seq enhancer activity with H3K27ac profiling gave us a catalog of enhancers with smaller region sizes compared to H3K27ac only (average region size: 1 vs 4 kb), with enhancers defined using two signals being more enriched in HaQTLs and TFs compared to those defined based on H3K27ac only. Taken collectively, our results suggest that the combination of CapSTARR and H3K27ac analyses might provide a more precise metric to describe enhancer functionality *in vivo*.

Genomic variants (including SNPs and SCNAs), and various TF binding enrichment at enhancer regions are two sources of enhancer heterogeneity. SNPs in enhancer regions may affect enhancer activity through several mechanisms, including alteration of TF binding behavior, chromatin accessibility change and alteration of H3K27ac levels [65]. Indeed, in our study, we observed a SNP (rs1464264) associated with H3K27ac levels of an enhancer and the expression of *ARL4C*. Notably, *ARL4C* has been reported to be a GC risk-associated gene identified in previous GWAS studies [66], and rs1464264 has been reported as an eQTL for *ARL4C* in Lymphoblastoid cells [67, 68]. This example deepens our knowledge of the functional impact of cancer risk-associated SNPs identified in GWAS studies.

Our study has limitations. First, our CapSTARR-seq data is based on only two cell lines. While our key findings were subsequently validated in multiple samples, we acknowledge that more CapSTARR-seq libraries will be required for provide a comprehensive overview of functional enhancer activity in GC. Second, we still lack a clear explanation of predicted enhancers that exhibit H3K27ac enrichment but no CapSTARR-seq activity. It is possible that the H3K27ac mark is not limited to enhancers, but may also mark other classes of distal regulatory elements targeted by CTCF [6, 17, 69]. We hypothesize that the CapSTARR-low/H3K27ac-high elements may represent insulators or other unknown regulators, which requires further study.

In conclusion, our study demonstrates that cell-line-predicted enhancers are pervasively linked to epigenomic and regulatory circuitry *in vivo*, and reveals mechanisms involving somatic CNVs, germline SNP variation, and *trans*-acting transcription factors in driving enhancer heterogeneity. We also provide evidence that combining histone modification and functional assay data provides a more accurate metric to assess enhancer activity than either platform individually, and identified novel genes associated with GC. Taken collectively, these studies will likely deepen our knowledge of enhancer functions driving GC development and progression.

## Methods

### Cell lines and primary tumor samples

Primary patient samples were obtained from the SingHealth Tissue Repository with approvals from institutional research ethics review committees and signed patient informed consent. ‘Normal’ (that is, nonmalignant) samples used in this study refer to samples harvested from the stomach, from sites distant from the tumour and exhibiting no visible evidence of tumour or intestinal metaplasia/dysplasia upon surgical assessment. Tumour samples were confirmed by cryosectioning to contain >40% tumour cells. AGS, Hs738, Hs746T, HsInt and SNU16 cells were obtained from the American Type Culture Collection. FU97, IM95, IST1, KATO-III, MKN7, NUGC4, OCUM1, RERF-GC-1B and SCH cell lines were obtained from the Japan Health Science Research Resource Bank. CLS145 and HGC27 cells were obtained from the Cell Lines Service. NCC19, NCC24, NCC59, SNU1750, SNU1967, SNU484 and SNU719 cell lines were obtained from the Korean Cell Line Bank. LMSU was obtained from the Riken Gene Bank. HFE145 cells were a gift from Dr. Hassan Ashktorab (Howard University, Washington, DC). GES1 cells were a gift from Dr. Alfred Cheng, Chinese University of Hong Kong. YCC3, YCC7, YCC10, YCC11, YCC21, and YCC22 were gifts from Yonsei Cancer Centre (Seoul, South Korea). MycoAlert

Mycoplasma Detection Kits (Lonza) and MycoSensor qPCR Assay Kits (Agilent Technologies) were used to detect mycoplasma contamination. All cell lines were negative for mycoplasma contamination.

### Nano ChIP-seq and data analysis

Nano-ChIP-seq was performed as described in the previously published protocol [29]. We assessed the qualities of ChIP-seq libraries (H3k27ac, H3K4me1 and H3K4me3) in two steps. First, we did quality control checks on the raw sequence reads generated from ChIP experiments through FastQC software (version 0.11.7). Second, we used two independent methods, CHANCE (CHip-seq ANalytics and Confidence Estimation) and ChIP enrichment assessment, to validate whether a library showed successful enrichment across the genome [29]. If proven high-quality, sequence reads were mapped to the human genome (hg19) using Burrows-Wheeler Aligner [70] (BWA-MEM, version 0.7.0), after trimming the first ten and last ten bases. Reads with high mapping quality (MAPQ) ≥ 10 retained for downstream analysis. To exclude the PCR amplification biases, fragments that had the same starting and ending coordinates were removed by the “rmdup” algorithm of Samtools (version 0.1.19). Peaks with significant ChIP enrichment relative to the input library (FDR<0.05) were detected using CCAT (version 3.0). Sequence coverage was computed using the R package “MEDIPS” (version 1.20.1) with a 50 bp bin size and read length extension to 200 bp. Peak density within a specific region was calculated by counting the total number of mapping reads normalized by the library size and the region length, an equivalence to the metric RPKM. To account for the background noise, background-corrected read densities were computed by subtracting the corresponding input signal from the ChIP signal.

### MeDIP-sequencing and data analysis

In brief, DNA was sonicated using COVARIS S2 and peak fragment distribution between 100-500bp was verified on an Agilent Bioanalyzer (Agilent Technologies) using the DNA1000 chip. Fragmented DNA was end-repaired, dA-tailed and adapter ligated using NEBNext® DNA Library Prep Master Mix Set for Illumina (E6040). Samples were then spiked with control DNAs that were unmethylated, methylated and hydroxymethylated (Diagenode C02040010) as a quality control measure. For each sample, input DNA that was not exposed to the primary antibody was included. Adapter-ligated DNA was subjected to immunoprecipitation with a primary monoclonal antibody against 5-methyl cytosine (Diagenode C15200081) using a previously published protocol [71]. Real-time PCR using primers against the spiked DNA controls were performed to verify successful and specific enrichment of methylated DNA. Immunoprecipitated samples were amplified using Phusion® High-Fidelity DNA Polymerase (M0530) and NEBNext® Multiplex Oligos for Illumina® (E7335) for 10 cycles. Amplified libraries were run on the Agilent Bioanalyzer using the High sensitivity DNA kit prior to Illumina sequencing using a single-end 100 base pair configuration. Reads were mapped to the human genome (hg19) using Burrows-Wheeler Aligner (BWA-MEM, version 0.7.0), after trimming the first ten and last ten bases. Reads with high mapping quality (MAPQ) ≥ 10 retained for downstream analysis. To exclude the PCR amplification biases, fragments that had the same starting and ending coordinates were removed by the “rmdup” algorithm of Samtools (version 0.1.19).

### Identification of predicted enhancers/super-enhancers

We detected the genomic regions enriched for H3K27ac ChIP signal, previously shown to mark the active cis-regulatory elements. To exclude potential promotor predictions, we selected the regions distant from known annotated TSSs (>2.5kb, GENCODE v19) as predicted enhancer elements. For each cell line, average profiling of histone modifications (H3K27ac, H3K4me1, and H3K4me3) and DNA methylation (5mC) on the predicted enhancers and annotated promoters (500bp flanking annotated TSSs) was plotted using the R package “ngsplot”. Predicted enhancer regions with at least one base overlap across multiple cell lines were merged using Bedtools (version 2.25.0) to form a consistent coordinate reference. Predicted enhancers were further subdivided into predicted super-enhancers or typical enhancers using the ROSE algorithm with the default parameters.

### *In vivo* validation of GC enhancer catalog

**“**Recurrent” predicted enhancers were identified as those enhancers occurring in at least two GC cell lines. For *t*-Distributed Stochastic Neighbor Embedding (*t*-SNE), we used signals from cell-line-predicted recurrent enhancers showing H3K27ac enrichment in two or more patients. Read densities over cell-line-predicted recurrent enhancers across GC samples and the matched normal tissues were corrected for potential batch effects using ComBat. *t*-SNE analysis was then performed using the R package “Rtsne” and plotted using R. WGS data from 212 GC samples was processed as described in the previously published paper [72]. The somatic point mutation rate over a set of regions was calculated as the total number of somatic point mutation calls divided by the sum of region lengths. Considering the variation in background mutation rates of GCs, the relative somatic point mutation rate was calculated as the log2 fold change of the somatic point mutation rate over the background point mutation rate.

### CapSTARR-seq and data analysis

CapSTARR-seq was performed as described in the previously published protocol[73]. Paired-end sequencing reads were aligned to human genome hg19 by BWA-mem. Uniquely mapped fragments with MAPQ ≥ 10 were collected for further analysis. To exclude the PCR amplification biases, fragments that had the same starting and ending coordinates were removed by the Samtools “rmdup” algorithm. To generate an overview of CapSTARR-seq signals, we created the circular visualization of CapSTARR-seq signals over regions within 2000 bases flanking probes across the human genome using the R package “RCircos” (version 1.2.1). To identify potential functional enhancers, we used macs2 (default setting; q value<0.05; keep-dup all; BAMPE mode; nomodel mode) to call peaks from all reads combined from three biological replicates for each cell line, OCUM1 or SNU16. We filtered out the enhancer peaks with their summits not overlapping CarStarr-seq probes or the corresponding predictive enhancer regions called by CCAT (using H3K27ac signals). The retained enhancers were defined as active functional enhancers and subsequently subdivided into four categories: inactive (fold change <1.5), weak (1.5 ≤ FC < 2), moderate (2 ≤ FC < 3) and strong (FC ≥ 3). Average profiles of histone modifications (H3K27ac, H3K4me1 and H3K4me3) and DNA methylation (5mC) over strong, moderate, or weak enhancers were generated by extracting background-corrected ChIP-seq signal from wiggle files around the summits of these selected enhancers. In detail, we calculated the mean RPKM values of epigenetics markers in each 50bp non-overlapping window spanning 6kb around the summits of enhancers.

We used Mann–Whitney U test to compare the CapSTARR-seq signals over four categories of predicted enhancers defined using H3K27ac signal. In detail, the CapSTARR-seq density within a special enhancer was computed using bigWigAverageOverBed (version 2). We calculated the number of TF binding sites over an enhancer using the ReMap database. To remove the potential confounding effect of DNA accessibility, DNA copy number and region length of predicted enhancers on CapSTARR-seq differentiation, we regressed out the effect of those three factors from CapSTARR-seq signal using a generalized linear model (GLM). In detail, DNA accessibility and DNA copy number of an enhancer is estimated using ATAC-seq data and WGS data, respectively. We compared the corrected CapSTARR-seq signals and TF binding enrichments over four categories of predicted enhancers using Mann–Whitney U test.

### ATAC-seq and data analysis

ATAC-seq was performed as previously described [73]. The reads were mapped to the human genome assembly (hg38) with Bowtie2. Mitochondrial and viral reads were filtered out. We called ATACseq peaks using MACS2 (callpeak function with parameters –nomodel and -B), and generated 300bp bins flanking each summit. We filtered promoter peaks (<2kb from TSS) and non-cell-specific peaks open in over 30% of ENCODE cell lines [74–76].

### ABC model

We selected all predicted enhancers captured by CapSTARR-seq probes as candidate regulatory elements in OCUM1 and SNU16 cells. Enhancer activity of candidate elements was assessed by using a combination of quantitative H3K27ac ChIP–seq and CapSTARR-seq signals. H3K27ac levels and enhancer strength assessed by functional enhancer assay are commonly used to predict enhancers and are predictive of the expression of nearby genes. Contact for each enhancer-gene pair in the ABC model was estimated using a function of the genomic distance between the enhancer and the TSS of the gene (Contact = Distance^-1^). 430 highly differentially expressed genes between OCUM1 and SNU16 cells were identified using a threshold method (Gene expression difference > 200 TPM). For each gene, all H3K27ac-defined enhancers within 5 Mb of the gene’s promoter were included as candidate functional enhancers. The total effect of all functional enhancers on target gene expression was assessed as the sum of ABC scores of those enhancers.

### GRO-cap and HiDRA data analysis

Mapping of transcription start sites (TSSs) captured by GRO-cap in human lymphoblastoid B cell (GM12878) and chronic myelogenous leukemic (K562) ENCODE Tier 1 cell lines was collected from the previously published paper [77]. H3K27ac enriched regions in GM12878 and K562 cells are available through the ENCODE data portal (www.encodeproject.org) under accession nos. ENCSR000AKC, ENCSR000AKP. To assess significance of enhancer-promoter interactions in the genomic regions containing GRO-cap TSS, we randomly permuted the positions of sequences within H3K27ac enriched regions distal to annotated promoters using shuffleBed with the -incl flag. To assess the significance of enrichment, we performed 10,000 shuffles of sequences. An empirical *P*-value was then derived by counting the number of times where the number of randomly shuffled sequences overlapping the enhancer regions interacting with promoters exceeded the observed number of genomic regions containing GRO-cap TSS overlapping the enhancer regions interacting with promoters. Raw sequencing files of the HiDRA dataset were obtained from the Sequence Read Archive (accession no. SRP118092) and processed as described by Wang et al [46]. By analyzing this dataset, we identified ∼47,000 promoter-distal genomic regions that are enriched for HiDRA signals which was referred to as “STARR active enhancers”. STARR-seq active regions in K562 cells were collected from the previous paper [78].

### RNA-seq analysis

RNA-seq was performed as described in the previously published protocol [79]. RNA-seq reads were mapped to the human reference genome (hg19) by using STAR (version 2.6.1a). The per-base sequencing quality and the per sequence quality scores of the mapped reads were assessed using FastQC. Transcript abundances at the gene level were calculated in the metric of TPM using RSEM (version 1.2.31).

### Identification of differential enhancers

CapSTARR-seq RPKM values were filtered by the predetermined log fold change (>1.5) and absolute difference (>5 RPKM) between OCUM1 and SNU16 cells. The same filtering was also performed on the H3K27ac ChIP data. As we have demonstrated that combining H3K27ac-enrichment and CapSTARR-seq signal to estimate enhancer activity outperforms the use of either of the features, 1,888 differential enhancers were identified from the union of regions obtained for CapSTARR and H3K27ac analyses.

### Identification of SNPs and CNAs

Sequencing reads from WGS libraries of GC cell lines were mapped to the human reference genome (hg19) using BWA. Duplicated reads marked by Picard (version 1.9.2) were removed. Indel regions were realigned by using GATK [80]. Base quality score recalibration was conducted by GATK. SNPs were called by using GATK Haplotype Caller. Copy number changes are generated by using CNVkit with the default parameters (bcbio-nextgen v0.9.3). As GC cell lines have no matched germline samples, CNVs were call against a non-matched normal sample. CNA breakpoints are defined as the ends of non-diploid segments. As the purity of cell lines is 100%, the DNA copy number of a segment is equal to 2^log2(tumor coverage/normal coverage). Copy number of an enhancer is defined as the copy number of the segment at which it is located.

### CNA-associated differential enhancer analysis

To detect differential enhancers associated with CNA, 996 differential enhancers with DNA copy number being abnormal in either SNU16 or OCUM1 were obtained (referred to as copy-number-abnormal enhancers thereafter). For each copy-number-abnormal enhancer, we employed the GLM model to test whether there is a significant correlation between H3K27ac signals and DNA copy numbers across 28 cell lines. At FDR threshold of 10%, 58 Copy-number-abnormal enhancers exhibiting a significant correlation between H3K27ac signals and DNA copy numbers were defined as Copy-number-associated enhancer. For each CN-associated enhancer, we used the Pearson correlation test in R package “stats” to estimate the significance of the correlation between expression levels of target genes and copy numbers across multiple cell lines. *P* values of the Pearson correlation tests were adjusted using the Benjamini-Hochberg method.

### SNP-associated differential enhancer analysis

To exclude the potential confounding effect of CNAs on identifying differential enhancers caused by SNPs, we focused on 892 differential enhancers located in diploid regions in both OCUM1 and SNU16 cells (referred to as copy-number-normal enhancers thereafter). As differential enhancers were defined by comparing OCUM1 and SNU16 cells, only SNP calls covered by non-reference bases in either of the two cells were retained. Finally, a set of 605 SNPs within copy-number-normal enhancers were obtained. We then employed the genotype-independent signal correlation and imbalance (G-SCI) pipeline to detect SNPs whose genotype correlates with histone acetylation levels (HaQTLs). To reduce experimental and other unknown variations, H3K27ac signal over differential enhancers across 28 cell lines were quantile-normalized before HaQTLs calling. For each SNP, the raw *P*-value obtained from the G-SCI test was adjusted using Benjamini-Hochberg method. At FDR threshold of 10%, 207 candidate HaQTLs were identified. To further explore how those HaQTLs cause histone acetylation alterations across cell lines, 207 HaQTLs were assessed for functional impact using RegulomeDB. 43 HaQTLs predicted by RegulomeDB [52] to influence protein DNA binding (RegulomeDB score 1 or 2) were retained for subsequent genotype-expression correlation analysis.

Gene expression profiles of 200 GC and 100 matched normal gastric samples from Singapore were generated using Affymetrix Human Genome U133 Plus 2.0 Array (GSE15459) and processed as described previously [30]. Genotype information of Singapore GC patients was extracted from the .CEL files produced from Affymetrix Genome-Wide Human SNP 6.0 Arrays (GSE31168) using the R package “crlmm”. Outliers classified as gene expression values that fall more than 1.5 interquartile ranges (IQRs) below the first quartile or above the third quartile. Outliers were excluded from each genotyping group to increase statistical power. *P* values were calculated using Student’s t-test.

### *ARL4C* shRNA knockdown and *ING1* overexpression

For *ARL4C* knockdown, shRNA sequences targeting *ARL4C* were cloned into pLKO.1 lentiviral plasmid vector and transfected into HEK293T for virus generation. shRNA sequences used were:shARL4C #1: CCGGGGAGCTGCGAAGTCTGATTTACTCGAGTAAATCAGACTTCGCAGCTCCTTTTTG shARL4C #2: CCGGCGAGGGCATGGACAAGCTCTACTCGAGTAGAGCTTGTCCATGCCCTCGTTTTTG shLuc: CCGGCACTCGGATATTTGATATGTGCTCGAGCACATATCAAATATCCGAGTGTTTTTG LMSU cells were infected with lentiviral particles and selected with 2ug/ml puromycin for at least 3 days before western blot validation and cell proliferation assays.

For *ING1* overexpression, the following primers were used to clone the *ING1* cDNA: ING1-F: CGACGATGACAAGGGATCCATGTTGAGTCCTGCCAACGGG ING1-R: GGAATTGATCCCGCTCGAGCTACCTGTTGTAAGCCCTCTC *ING1* cDNA was cloned into a pHR‟CMVGFPIRESWSIn18 based vector (gift from Dr. Shang Li, Duke-NUS) using Gibson Assembly and transfected into HEK293T for virus generation. LMSU cells were infected with lentiviral particles and selected with 400ug/ml hygromycin for at least 3 days before western blot validation and cell proliferation assays.

### Western blot

Cells were harvested and lysed in RIPA buffer (Sigma) and incubated on ice for 10 min. Lysates were cleared by centrifuging at 9000 rpm for 10 min. Protein concentrations were measured using the Pierce BCA protein assay kit (Thermo Scientific). Samples were diluted in 4X Laemmli Buffer (Biorad), boiled at 95°C for 10 min and loaded for SDS-PAGE. The following antibodies were used: ARL4C (#10202-1-AP, Proteintech), ING1 (#14625, Cell Signalling Technology) and GAPDH (#60004-1-Ig, Proteintech).

### Cell proliferation assays

Cell proliferation rates were measured using the Cell Counting Kit-8 (Dojindo). Briefly, 1000-3000 cells were seeded in 96-well plates and cell density was measured every 2 days following manufacturer’s protocol. *P*-values were derived using Student’s t-test with *P*-values <0.05 considered as statistically significant.

### Survival analysis

300 GC patients in the ACRG cohort were collected to study the association between expression levels of selected genes and GC survival. We employed Kaplan–Meier survival analysis with the overall survival as the outcome metric. Log-rank tests were used to estimate the significance of Kaplan– Meier curves. Gene expression of the ACRG cohort was profiled using Affymetrix Human Genome U133plus 2.0 Array (GSE62254 and GSE66222) and processed as described previously [28].

### TF-associated differential enhancer analysis

We interrogated enrichments of TFs in OCUM1-specific enhancers and SNU16-specific enhancers using HOMER with default parameters. The top 5 TF identified from the HOMER outputs were used for differential expression analysis. Level 3 TCGA RNA-seq normalized matrix for 415 GC and 35 normal gastric samples was downloaded from the Broad Institute TCGA Genome Data Analysis Center (GDAC) Firehose (https://gdac.broadinstitute.org/). HNF4α-associated differential enhancers were defined as those SNU16-specific enhancers exhibiting a significant correlation between HNF4α binding enrichment and H3K27ac signals among multiple lines. HNF4α ChIP data of GC cell lines and RNA-seq data of HNF4α perturbation were downloaded from GSE114018 and were processed as described previously [79].

## Supplementary information

Additional file 1: Supplementary Figures and Figure legends S1-S15. Additional file 2: Supplementary Tables S1-S5.

## Acknowledgements

We thank the Sequencing and Scientific Computing teams at the Genome Institute of Singapore for sequencing services and data management capabilities, and the Duke-NUS Genome Biology Facility for sequencing services. We also thank Liu Mo, Steve Rozen, Melissa Jane Fullwood and Heike Irmgard Grabsch for helpful discussions.

## Author contributions

S.W.T.H., C.X., M.X., N.P., L.M., M.R., C.G.A.N. and R.S.Y.F. performed the experiments. T.S., S.W.T.H., W.F.O., K.K.H., Y.A.G., S.N.L., M.M.C., M.R.M. and M.A.B. analyzed the data. A.L.K.T., X.O. and A.J.S. provided facilities, reagents, and intellectual input. P.T., S.J. and K.P.W. supervised the study. T.S., S.W.T.H., and P.T. wrote the paper.

## Availability of data and materials

All raw and processed sequencing data generated in this study have been submitted to the NCBI Gene Expression Omnibus (GEO; https://www.ncbi.nlm.nih.gov/geo/) under accession number GSE162420. Previously deposited histone ChIP-seq (GSE51776, GSE76153, GSE75898 and GSE121140) and RNA-seq (GSE85465, GSE121140) data that are used in this study are available in Gene Expression Omnibus. CapSTARR-Seq and ATAC-seq data sets are available from Gene Expression Omnibus under GSE121140. Other public data sets used are described in the Method part.

## Ethics approval and consent to participate

This study did not require any ethics approval.

