## Supplementary Figures for "Integrative Epigenomic and High-Throughput Functional Enhancer Profiling Reveals Determinants of Enhancer Heterogeneity in Gastric Cancer"

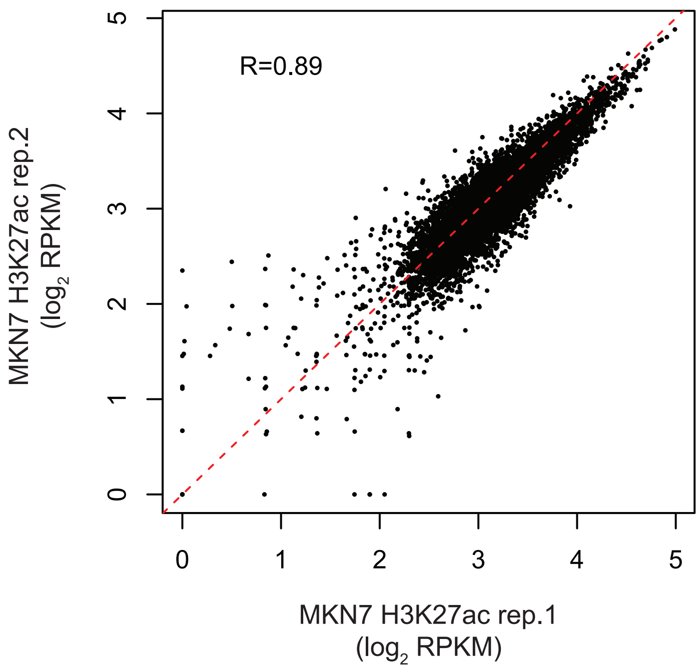


**Supplementary Figure 1: Comparison of H3K27ac signals over common predicted enhancers between two MKN7 replicates.**

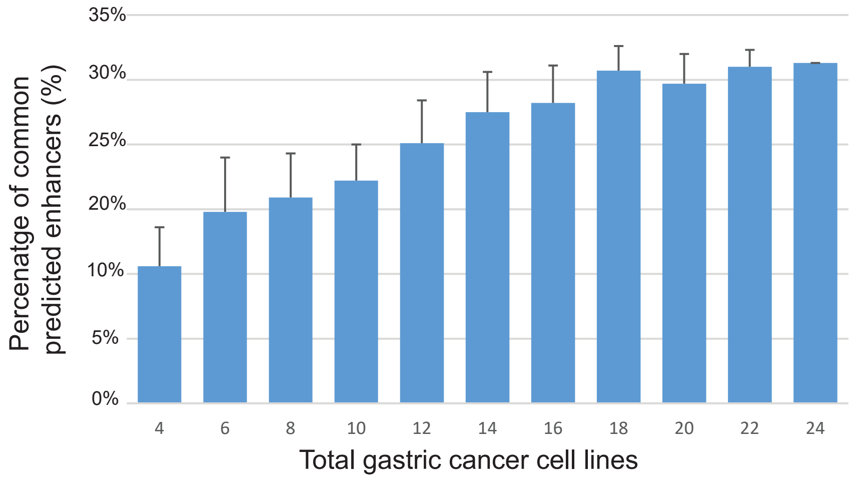


**Supplementary Figure 2: Recurrence rates of predicted enhancers.** Recurrent enhancers here are defiend as those enhancers occuring in at least 3 GC cell lines Data presented are the mean percentage +/- standard deviation of common predicted enhancers found in two or more gastric cancer cell lines, as a function of number of cell lines.

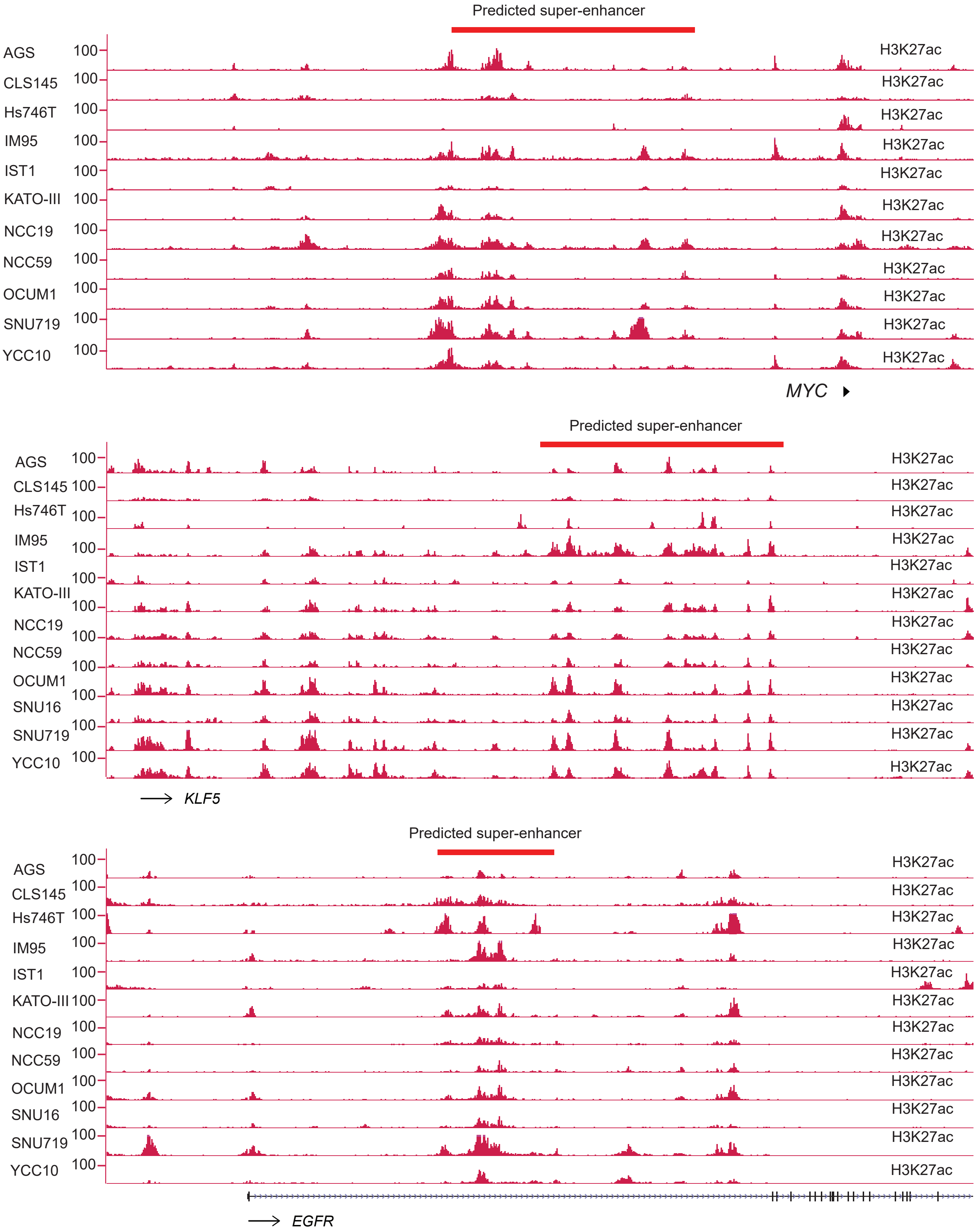


**Supplementary Figure 3: Predicted super-enhancers in GC cell lines.** *MYC*-, *KLF5* and *EGFR*-associated predicted super-enhancers are recurrent in multiple GC cell lines.

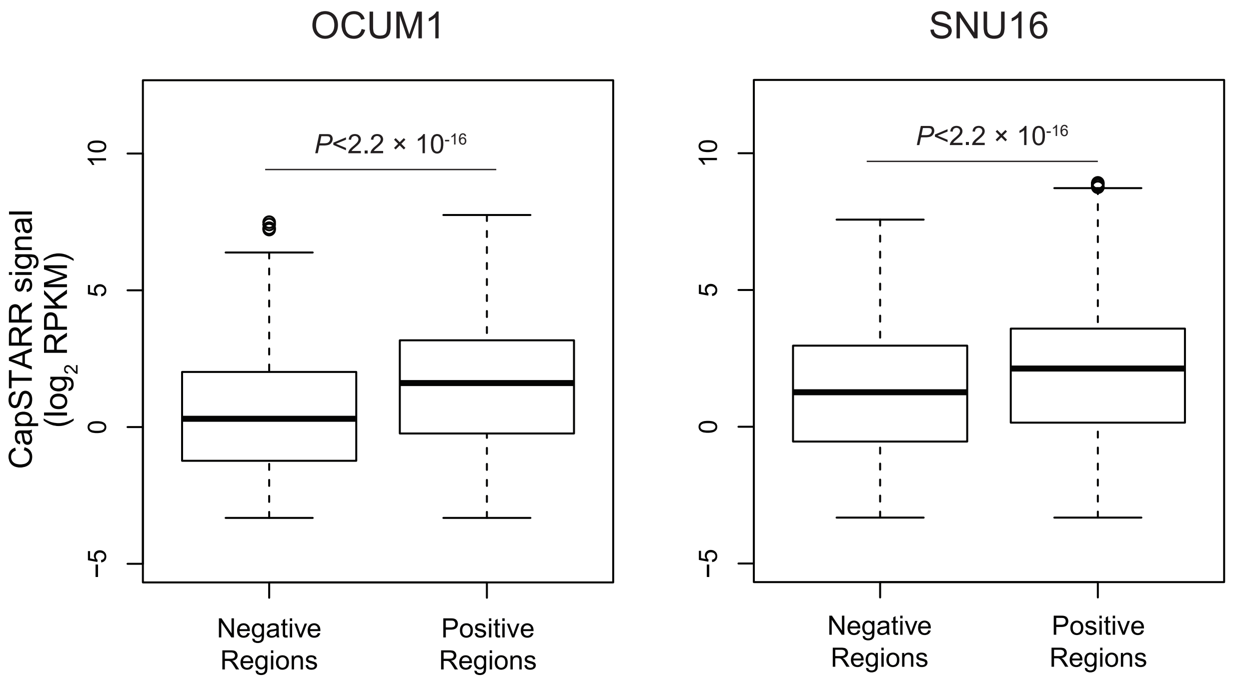


**Supplementary Figure 4: Differences in corrected CapSTARR-seq signals ($\log_2$ RPKM) over captured predictive enhancers (positive) and negative regions in OCUM1 and SNU16 cells.** *P*-values: Mann–Whitney U test.

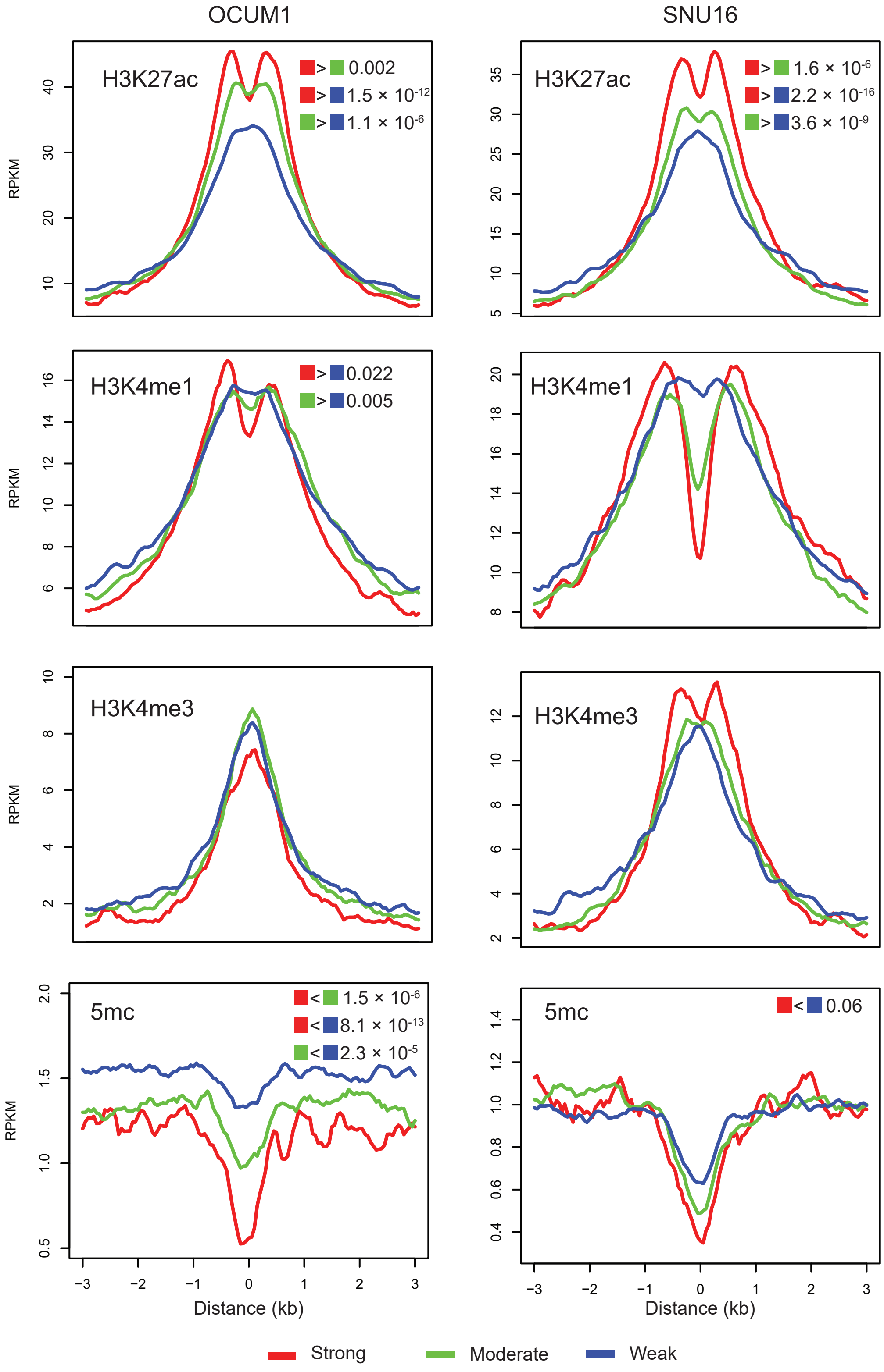


**Supplementary Figure 5: Average profiles of H3K27ac, H3K4me1, H3K4me3 ChIP-seq and DNA methylation (5mC) for CapSTARR-seq enriched regions (CERs) classified as weak, moderate or strong in OCUM1 (left panels) or in SNU16 (right panels).** Statistically significant differences calculated on the regions within 1k bases flanking the summits of CERs.

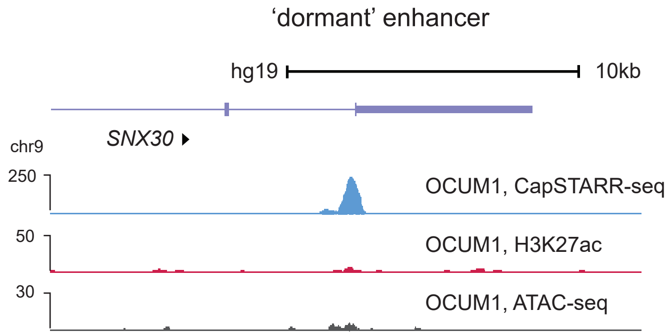


**Supplementary Figure 6: CapSTARR-seq (blue), H3K27ac ChIP-seq (red) and ATAC-seq (black) tracks at SNX30 locus in OCUM1 cells.** An OCUM1 ‘dormant’ enhancer exhibits high CapSTARR-seq signals but depletion of H3K27ac and ATAC-seq peaks.

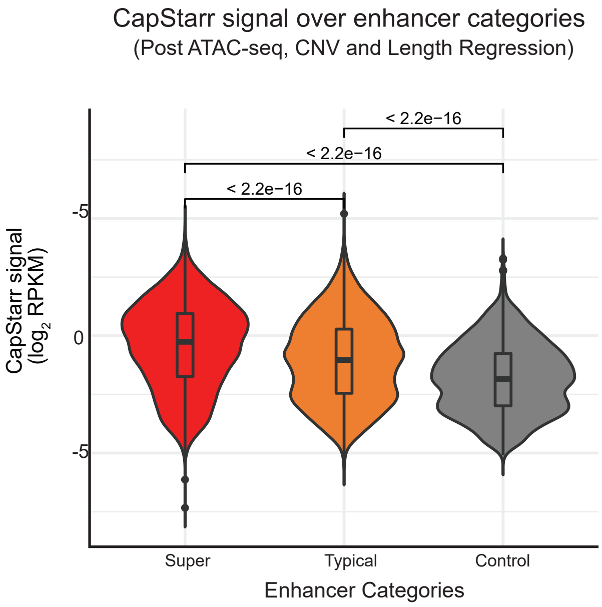


**Supplementary Figure 7: Differences in corrected CapSTARR-seq signals (log2 RPKM) in enhancer categories in OCUM1 cells.** Effects of DNA accessibility, DNA copy number and region length of predicted enhancers were regressed out from Capstarr-seq signal using a generalized linear model (GLM). *P*-values are calculated using the Mann–Whitney U test.

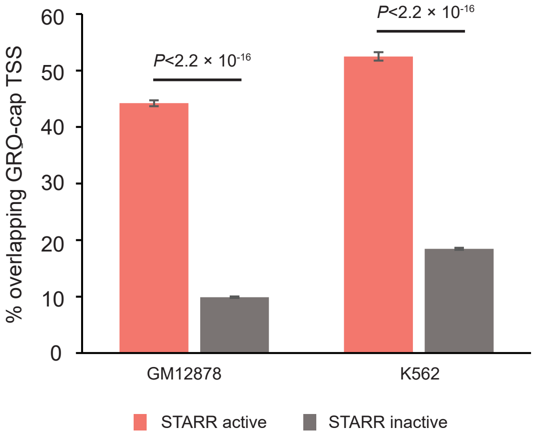


**Supplementary Figure 8: The percentage of STARR active and inactive elements within H3K27ac active enhancers overlapping GRO-cap TSS.** The error bars indicate the s.e.m. calculated for a sample of binary trials, centered on the observed success rate. P values are from a two-sided Fisher’s exact test.

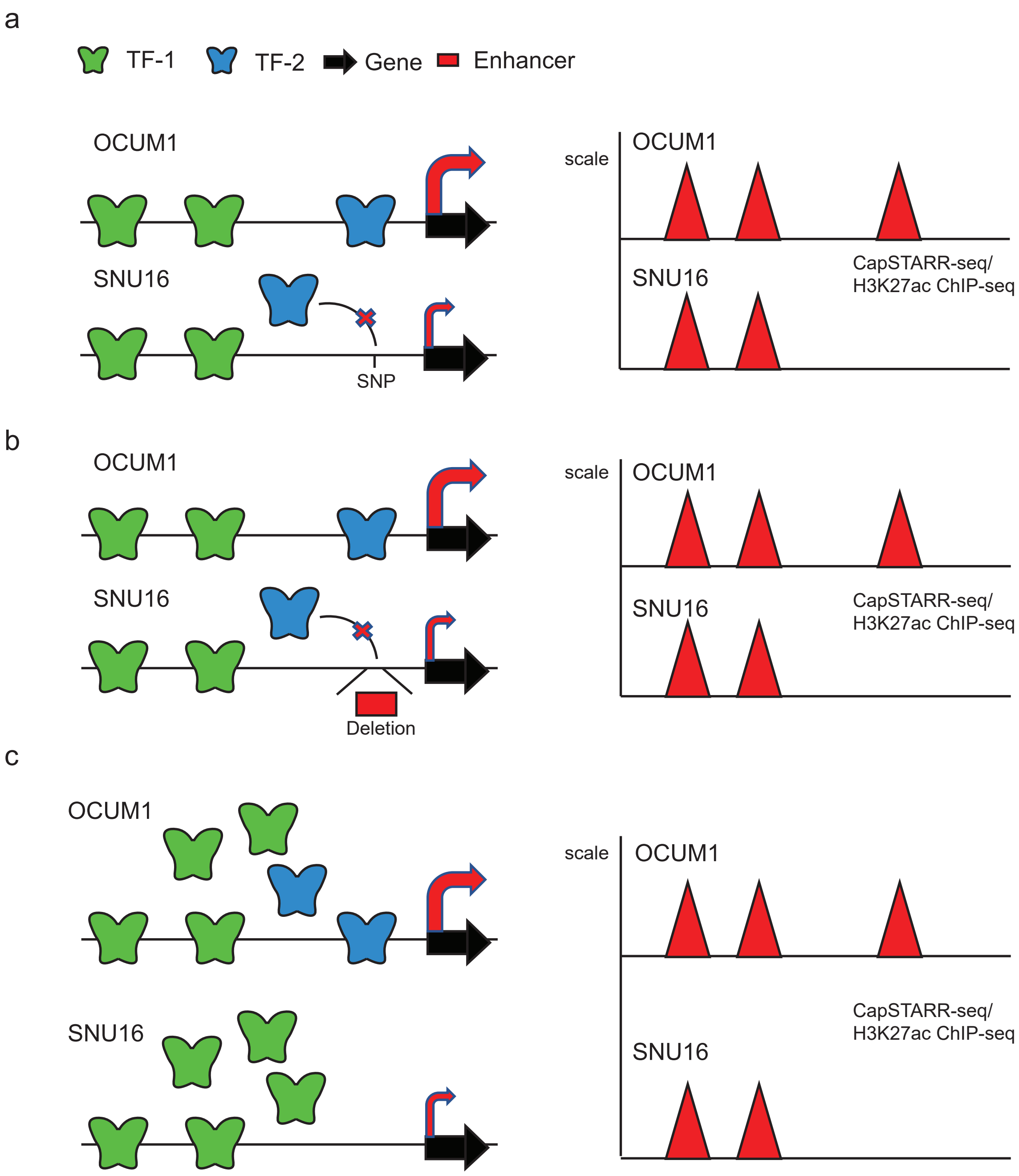


**Supplementary Figure 9: Schematic diagram of '*cis*-model' (panel a, b) and '*trans*-model' (panel c).**
**a)** Existence of some SNP causes differential TF binding enrichment, resulting in differential CapSTARR-seq/H3K27ac ChIP-seq signals over the enhancer.
**b)** Copy number variation of the enhancer region causes differential TF binding enrichment, resulting in differential CapSTARR-seq/H3K27ac ChIP-seq signals over the enhancer.
**c)** Low expression of some TF results in the disappearance of the CapSTARR-seq/H3K27ac ChIP-seq signal over the enhancer region.

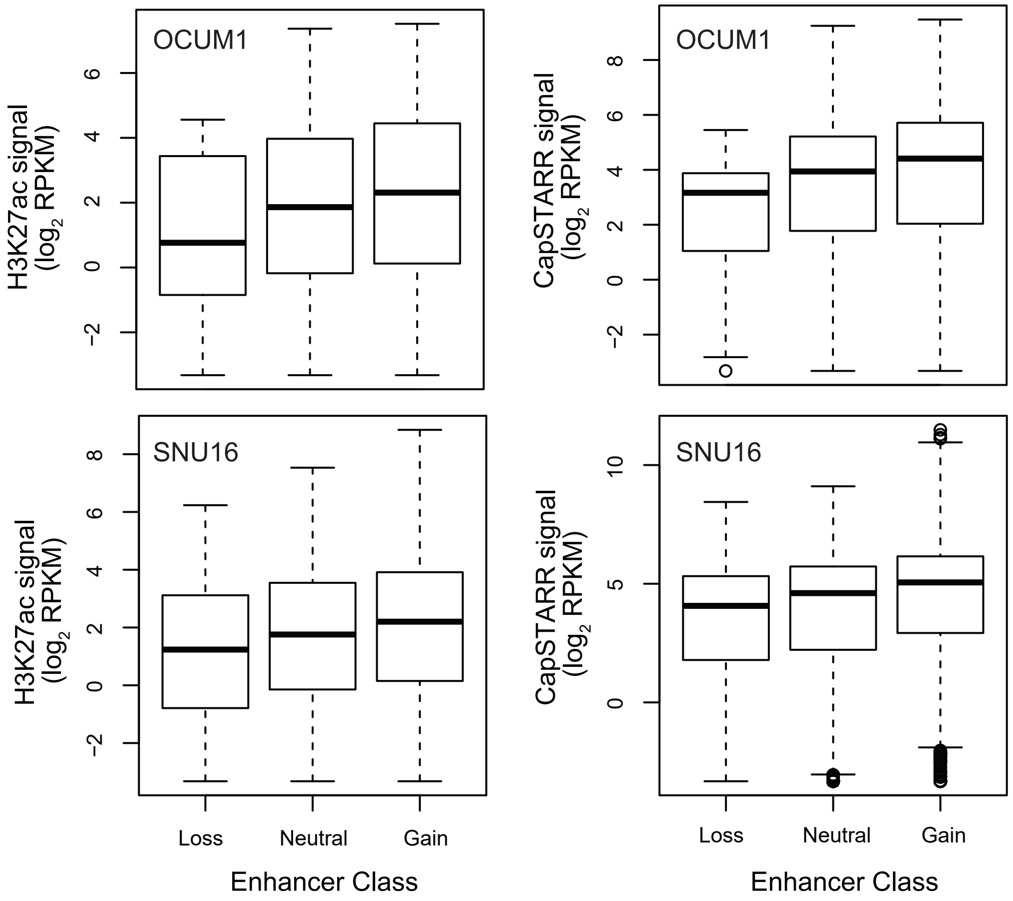


**Supplementary Figure 10: Differences in H3K27ac and CapSTARR-seq singals ($log_2$ RPKM) among three enhancer classes (Loss, Neutral and Gain).** Loss enhancers are those with DNA copy number smaller than 2. Neutral enhancers are those with DNA copy number equal to 2. Gain enhancers are those with DNA copy number larger than 2.

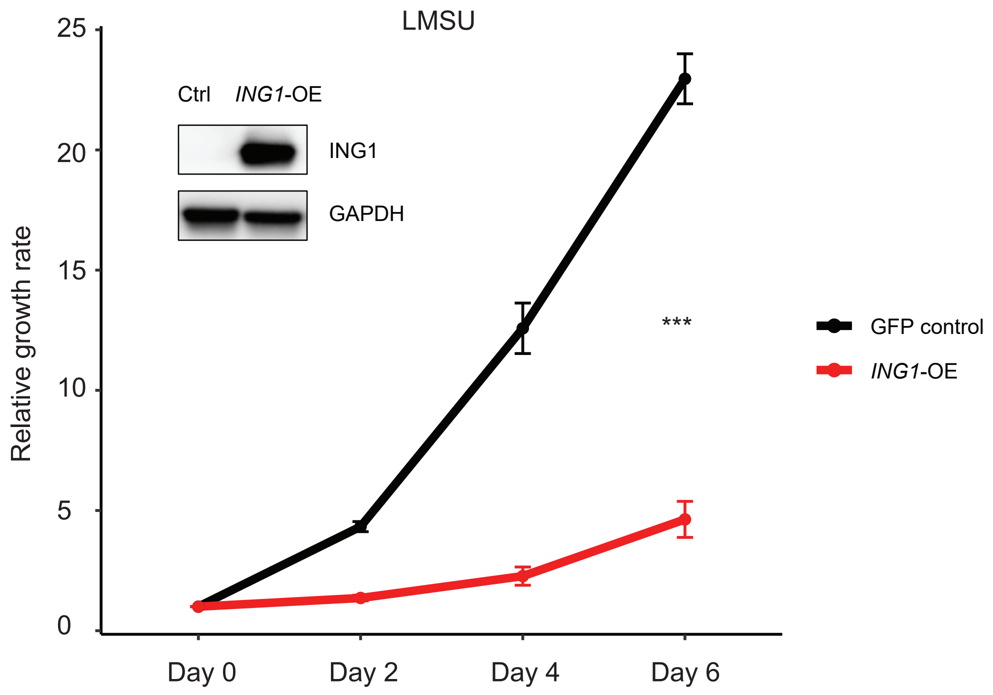


**Supplementary Figure 11: Relative cell proliferation rate after *ING1* overexpression in LMSU.** *ING1* overexpression led to a decrease in cell proliferation capacity. *$P$ < 0.05, **$P$ < 0.01, and ***$P$ < 0.001 by 2-sided Student's t-test. Error bars indicate the SD. All data are representative of 3 independent experiments.

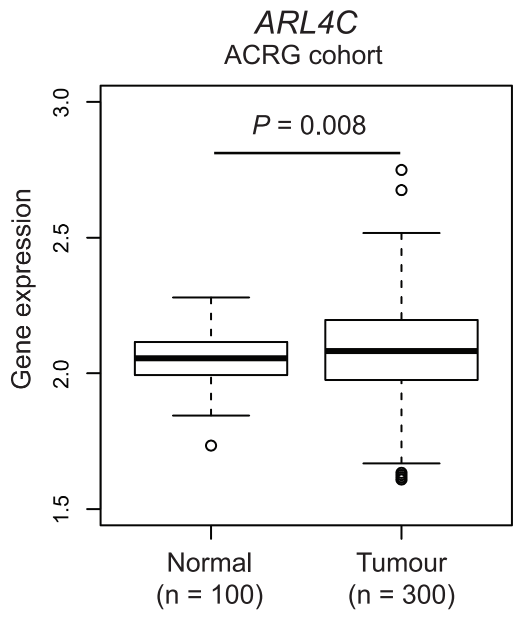


**Supplementary Figure 12: Expression of *ARL4C* in normal gastric (n=100) and GC samples (n=300) from the ACRG cohort.** *P*-value: Student's t-test.

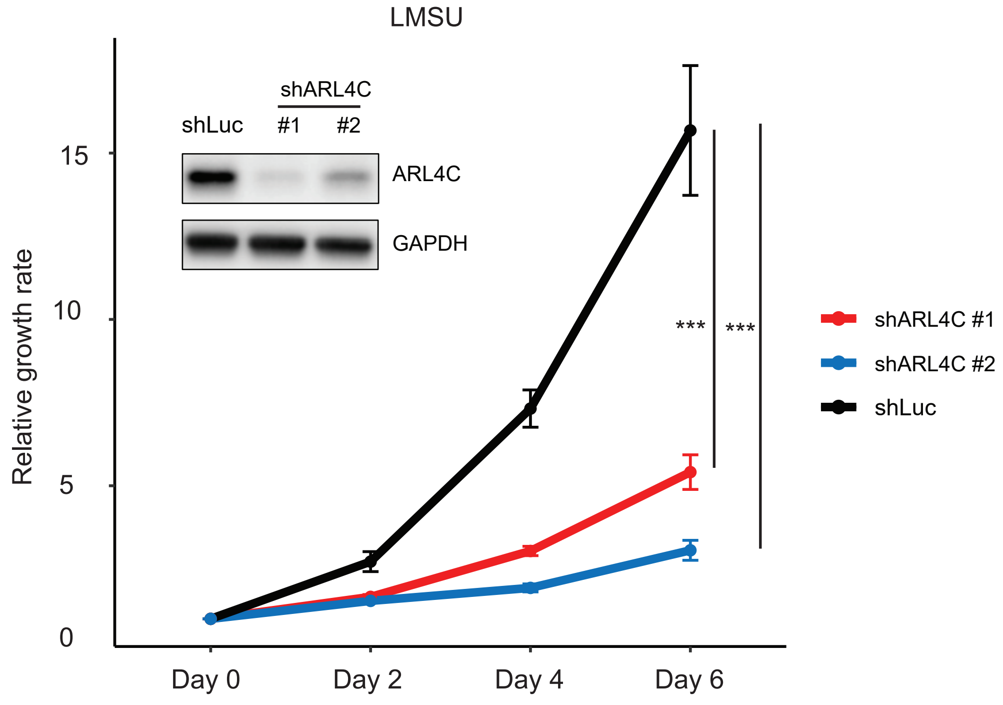


**Supplementary Figure 13: Relative cell proliferation rate after *ARL4C* knockdown in LMSU.** *ARL4C* knockdown led to a decrease in cell proliferation capacity. *$P < 0.05$, **$P < 0.01$, and ***$P < 0.001$ by 2-sided Student's t-test. Error bars indicate the SD. All data are representative of 3 independent experiments.

a

| Transcription factor | Motif | *P*-value | % Target regions with motif |
|---|---|---|---|
| Fra1 | 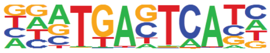 | $1\times 10^{-166}$ | 32.08% |
| Fra2 | 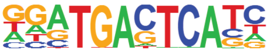 | $1\times 10^{-163}$ | 29.97% |
| Atf3 | 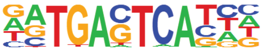 | $1\times 10^{-155}$ | 34.01% |
| BATF | 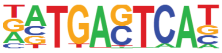 | $1\times 10^{-154}$ | 33.62% |
| JunB | 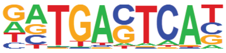 | $1\times 10^{-153}$ | 31.03% |

b

| Transcription factor | Motif | *P*-value | % Target regions with motif |
|---|---|---|---|
| Fosl2 | 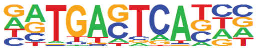 | $1\times 10^{-79}$ | 9.93% |
| BATF | 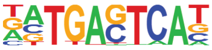 | $1\times 10^{-78}$ | 15.57% |
| Jun-AP1 | 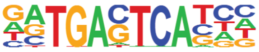 | $1\times 10^{-77}$ | 8.16% |
| Fra2 | 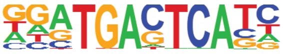 | $1\times 10^{-76}$ | 12.56% |
| Fra1 | 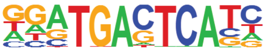 | $1\times 10^{-75}$ | 13.83% |
| Atf3 | 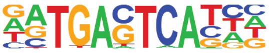 | $1\times 10^{-69}$ | 15.15% |

**Supplementary Figure 14: Top 5 transcription factor binding enrichments at OCUM1-specific enhancers defined based on a) CapSTARR and H3K27a paterns or b) only H3K27ac as determined by using HOMER *de novo* motif analysis.**

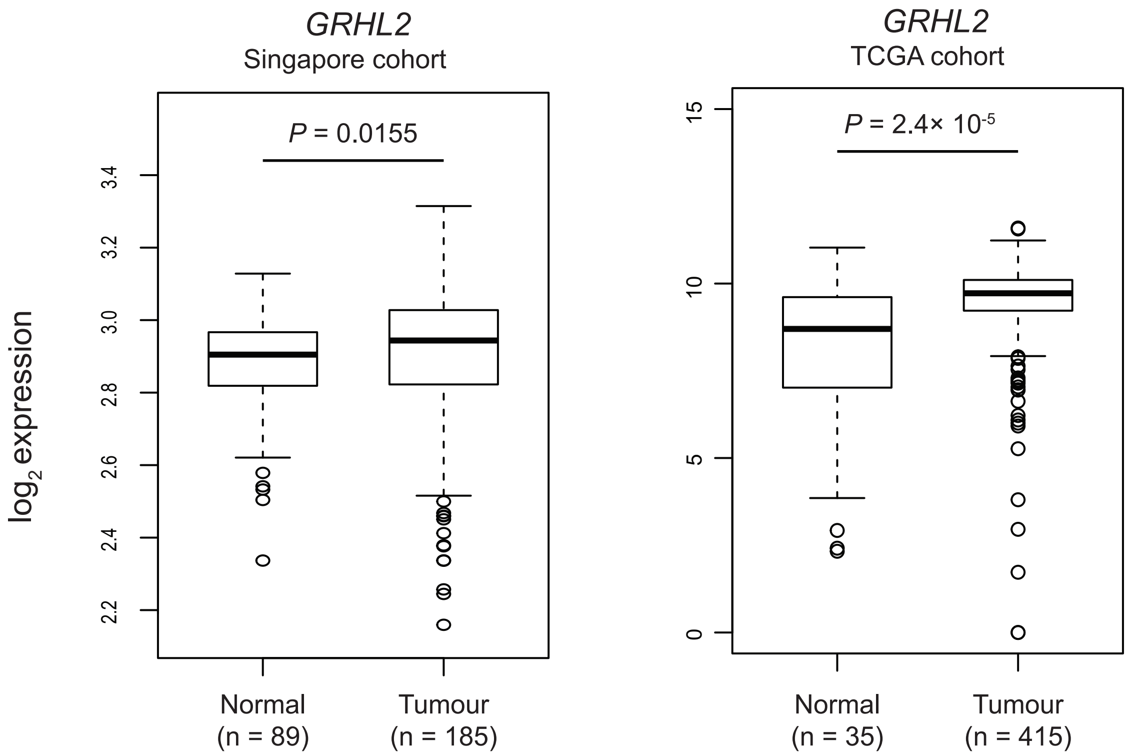


**Supplementary Figure 15: Expression of *GRHL2* in normal gastric and GC samples from the Singapore cohort (Left) or the TCGA cohort (Right).** *P*-value: Mann–Whitney U test.
